# Decoding stage-specific symbiotic programs in the *Rhizophagus irregularis*–tomato interaction using single-nucleus transcriptomics

**DOI:** 10.64898/2026.01.22.701092

**Authors:** Naomi Stuer, Toon Leroy, Thomas Eekhout, Annick De Keyser, Jasper Staut, Bert De Rybel, Klaas Vandepoele, Petra Van Damme, Judith Van Dingenen, Sofie Goormachtig

## Abstract

Arbuscular mycorrhizal fungi (AMF) establish a dynamic and asynchronous symbiosis with a wide range of land plants, involving distinct stages of root colonization and associated cellular responses that co-occur within the same root. Whilst decades of research have significantly advanced our understanding of the plant’s symbiotic gene repertoire, this spatial and temporal complexity has hindered a detailed dissection of the molecular mechanisms underlying fungal accommodation. Here, we present the first single-nucleus RNA-sequencing (snRNA-seq) dataset of *Solanum lycopersicum* roots colonized by *Rhizophagus irregularis*. Unsupervised subclustering of an AM-specific cell population resolves AM-responsive root epidermal cells as well as a developmental gradient of cortical cells across distinct stages of arbuscule formation, unveiling stage-specific transcriptional signatures during AMF colonization. Moreover, using Motif-Informed Network Inference based on single-cell EXpression data (MINI-EX), we put forward candidate transcription factors orchestrating these stage-specific transcriptional programs. Together, our data support novel hypotheses on how diverse plant developmental and physiological processes – including localized cell cycle reactivation and the integration of multiple nutritional cues – are coordinated to facilitate the establishment of a functional symbiosis. As such, this high-resolution dataset serves as a valuable resource for candidate gene prioritization and future reverse genetic studies.

## INTRODUCTION

The symbiotic relationship between arbuscular mycorrhizal fungi (AMF) and land plants is one of the oldest and most widespread mutualisms in the plant kingdom.^1-3^ In exchange for photosynthetically derived carbon (C), mainly supplied as lipids, AMF provide their host plants with improved access to nutrients and enhanced resistance to a wide range of biotic and abiotic stresses.^4-6^ Over the past four decades, extensive research has significantly advanced our understanding of the plant symbiotic gene repertoire.^1,7^ However, the precise spatiotemporal expression patterns of these genes, and regulation thereof, during arbuscular mycorrhiza (AM) symbiosis remains poorly understood.

Deciphering the molecular interplay that facilitates AM symbiosis is complicated by the asynchronous nature of fungal colonization: within a single root system, various interaction stages occur simultaneously among the diverse root cell types, each expected to display unique transcriptional responses to AMF colonization. These stages range from epidermal cells involved in early signaling and hyphopodium contact, to cortical cells interacting with intra- and intercellular hyphae, and inner cortical cells containing arbuscules at various developmental stages – from arbuscule initiation and active nutrient exchange to degradation.^1,7^ Moreover, even non-colonized cells exhibit symbiosis-related transcriptional changes, reflecting shifts in root metabolism, nutrient allocation, and systemic alterations in (a)biotic stress resilience.^8-10^ Traditional bulk transcriptomic approaches, which average gene expression levels across heterogeneous cell populations, obscure this cellular and developmental resolution. As a result, key regulatory mechanisms – such as those governing hormone signaling, immune modulation, metabolism, and nutrient transport – remain difficult to disentangle. This complexity highlights the need for high-resolution, stage-specific investigations into the host transcriptional programs that support AMF colonization.

The asynchronous nature of AM symbiosis has previously been addressed using laser-capture microdissection (LCM) to isolate arbuscule-containing cells. In legumes, such as *Medicago truncatula*^9,11-14^ and *Lotus japonicus*,^15-17^ and in the non-legume *Solanum lycopersicum* (tomato),^18,19^ LCM enabled profiling of symbiosis-related gene expression at an enhanced resolution. However, drawbacks of this approach include its labor-intensive nature, the need for manual identification of colonized cells, which may introduce sampling bias, and susceptibility to contamination from adjacent tissues.^20^

Single-cell and single-nucleus RNA-sequencing (scRNA-seq and snRNA-seq) circumvent these limitations by enabling the high-throughput, unbiased profiling of tens of thousands of individual cells or nuclei.^20-22^ Thanks to their unbiased nature and high resolution, single-cell transcriptomic methods enable the detection of rare or transient symbiotic states. Recent applications of sc/snRNA-seq in root nodule symbioses – including in *M. truncatula*, *L. japonicus*, and *Glycine max* – have demonstrated their power to resolve transcriptional heterogeneity in root–microbe interactions.^23-27^ Yet, their application to AM symbiosis remains limited to the study by Serrano et al., who examined the *M. truncatula*–*Rhizophagus irregularis* interaction by combining snRNA-seq with spatial transcriptomics, suggesting transcriptional signatures associated with different stages of colonization within the cortex.^28^

Expanding the application of snRNA-seq to study AM symbiosis in additional host species is critical, given that while a subset of symbiotic genes may be commonly induced across diverse plant species, AM-induced molecular pathways and their regulation can vary substantially between taxa.^29,30^ Here, we present an snRNA-seq analysis of *R. irregularis*-colonized roots of tomato, the most widely grown vegetable crop after *S. tuberosum* (potato), another member of the *Solanaceae*. By enriching for colonized root regions using *GFP* expression driven by the promoter of the arbuscule-induced tomato *PHOSPHATE TRANSPORTER 4* (*pSlPT4*),^31^ we were able to transcriptionally profile a large population of colonized cells representing major AMF colonization stages. We elaborate on the specific transcriptional fingerprints associated with colonization of epidermal cells, as well as with arbuscule development and maturation. Moreover, using Motif-Informed Network Inference based on single-cell EXpression data (MINI-EX), we predicted transcription factors (TFs) regulating colonization progression and efficiency at distinct stages. Not only does this systems-level approach aid in prioritizing candidate TFs involved in arbuscule development for future research, it also enhances our understanding of how diverse environmental and developmental cues are integrated to ensure a productive symbiosis.

## RESULTS

### Establishment and annotation of an snRNA-seq dataset of tomato–AM symbiosis

AM symbiosis is characterized by its asynchronous colonization dynamics, with distinct stages of the interaction co-occuring within the same root system.^1^ To gain an in-depth understanding of cell type- and interaction stage-specific transcriptional profiles during the tomato–*R. irregularis* symbiosis, snRNA-seq was performed on mock- and AMF-inoculated tomato roots, harvested at 6 weeks post inoculation (wpi) (Figures 1A-1E).

**Figure 1.**
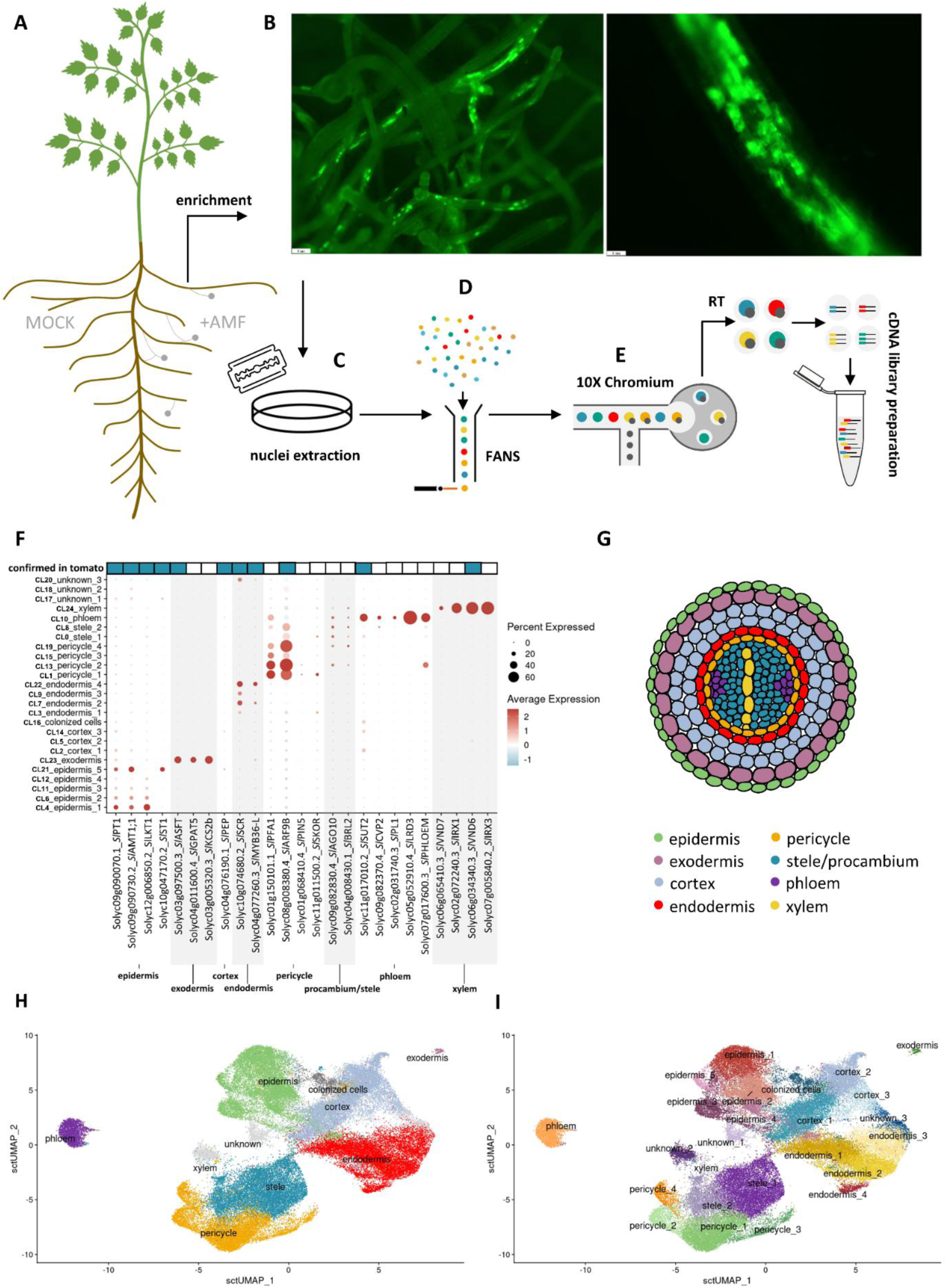
Overview of the workflow used for snRNA-seq of *R. irregularis*-colonized tomato roots and dataset annotation. **A.** Composite tomato plants with roots expressing *pSlPT4:GFP* were generated and grown in the presence (AMF) or absence (MOCK) of *R. irregularis*. Roots were harvested at 6 weeks post inoculation (wpi). **B.** *pSlPT4:GFP* promoter activity visualized using the Leica M165 FC fluorescence bino. Scale bars = 1mm. For AMF samples, only *GFP*-expressing root regions were selected, enriching for root tissues actively colonized by AMF. **C.** Nuclei were isolated from harvested roots using the chopping method on fresh tissue. **D.** Isolated nuclei were stained with propidium iodide (PI) and purified from debris via fluorescence assisted nuclei sorting (FANS). **E.** Schematic overview of snRNA-seq using the 10x Chromium platform. Within a microfluidic chamber, nuclei are co-encapsulated with gel beads containing barcoded oligonucleotides. After reverse transcription (RT) and cDNA synthesis, a cDNA library is generated, in which transcripts from individual nuclei are tagged with unique molecular identifiers. **F.** Expression profiles of 25 cell type-specific marker genes across all identified clusters (Table S1). Dot size indicates the percentage of nuclei within each cluster expressing the marker gene. Dot color represents the average scaled expression level, with red indicating higher and blue lower normalized expression. The top row indicates whether (blue) or not (white) marker specificity had previously been confirmed in tomato. **G.** Schematic representation of the various root cell types in a transverse section of a tomato root. **H.** UMAP plot depicting cluster assignment to major root tissue types (color-coded as in G). **I.** UMAP plot depicting all identified clusters and subpopulations.

To ensure analysis of highly colonized tissues, snRNA-seq was performed on composite plants with hairy roots harboring the *pSlPT4:GFP* construct, in which *GFP* is driven by the *PHOSPHATE TRANSPORTER 4* (*SlPT4*; Solyc06g051850.2) promoter, a marker for mature arbusculated cells. Only root sections displaying abundant GFP fluorescence were harvested, ensuring active fungal colonization in the AMF-colonized samples (Figure 1B). The activation of the *pSlPT4* promoter in tomato arbusculated cells was confirmed in hairy roots expressing *pSlPT4:GUS* in conjunction with WGA-AF488 staining for visualization of fungal hyphae (Figure S1). Plant nuclei were isolated, stained with propidium iodide, and purified by fluorescence-activated nuclei sorting to include all major root tissue types of AMF-inoculated and mock samples. Sorted nuclei were subsequently loaded onto a microfluidic chip for snRNA-seq profiling using the 10X Genomics Chromium platform (Figures 1C-1E).

For each condition (mock and AMF), snRNA-seq datasets were generated from three independent biological replicates, each comprising transcriptomes of approximately 10,000 nuclei (Supplementary file S1). The individual datasets were merged into a final Seurat object comprising 65,878 high-quality nuclei – 35,357 from mock-inoculated and 30,521 from AMF-inoculated samples. To correct for potential batch effects, Harmony was applied,^32^ resulting in improved integration across datasets, as evidenced by more cohesive clustering of nuclei in the Uniform Manifold Approximation and Projection (UMAP) embedding (Figure S2A). Unsupervised clustering identified 25 distinct clusters (Figure S2B), most of which were represented across all biological replicates and both treatment conditions. However, cluster 19, 22, and especially cluster 18, were predominantly derived from the first biological replicate (MOCK1 and AMF1) (Figure S2C). Further inspection of transcript and gene counts per nucleus within each cluster revealed that nuclei in cluster 18 exhibited a relatively low transcript abundance, suggesting that this cluster likely represents a technical artefact, rather than a true biological signal (Figure S3). Indeed, technical variation may have arisen from the use of different chemistries: the first replicate was processed using 10X Genomics NextGEM 3’ v3.1, while the second and third replicates used 10X Genomics GEM-X 3’ v4 (Supplementary file S1).

Cell cluster annotation was performed by analyzing the expression profiles of known marker genes for major root tissue types (epidermis, exodermis, cortex, endodermis, pericycle, phloem, procambium/stele, and xylem) across the distinct clusters of the snRNA-seq dataset (Figures 1F, 1G, and S4). The following genes exhibit tissue-specific expression in *S. lycopersicum*: the epidermis markers *PHOSPHATE TRANSPORTER 1* (*SlPT1*; Solyc09g090070.1), *AMMONIUM TRANSPORTER 1* (*SlAMT1*; Solyc09g090730.2), the potassium (K^+^) channel *SlLKT1* (Solyc12g006850.2)^33^ and *SULFATE TRANSPORTER 1* (*SlST1*; Solyc10g047170.2); the exodermis marker *SlASFT* (Solyc03g097500.3), a *SUBERIN FERULOYL TRANSFERASE^34^*; the cortex marker *SlPEP* (Solyc04g076190.1), a *PEPSIN-TYPE PROTEASE^35^*; the endodermis markers *SCARECROW* (*SlSCR*; Solyc10g074680.2)^35^ and the MYB TF *SlMYB36-L* (Solyc04g077260.3)^36^; the pericycle marker *AUXIN RESPONSE FACTOR 9B* (*SlARF9B*; Solyc08g008380.4)^37^; the phloem marker *SUCROSE TRANSPORTER 2* (*SlSUT2*; Solyc11g017010.2)^38^; and the xylem marker *VASICULAR-RELATED NAC-DOMAIN* (*SlVND6*; Solyc06g034340.3)^35^ (Figure 1F). Additional markers were selected based on their homology to well-characterized marker genes in other plant species (Table S1). Using this approach, we successfully annotated the majority of clusters and subpopulations to their corresponding major root tissue types (Figure 1H), often grouping multiple diverse clusters present in the original UMAP (Figure 1I).

To assess the accuracy of our cell type annotations, we compared them with annotations from a previously published tomato root atlas generated using scRNA-seq by Cantó-Pastor et al.^34^ To this end, we transferred labels from the reference scRNA-seq dataset using Seurat’s anchor-based label transfer. For several clusters, the transferred labels showed strong concordance with the original annotations, with high prediction scores indicating confident assignments (Figure S5). Prediction scores were particularly high for clusters corresponding to the stele (including procambium, phloem, and xylem), cortex, and pericycle. In contrast, only subsets of the epidermis and endodermis could be reliably annotated using the scRNA-seq reference.^34^ These discrepancies are likely attributable to differences in experimental methodology (single-cell vs. single-nucleus RNA-seq), developmental stage (1 week vs. 10 weeks post germination), plant growth conditions (*in vitro* vs. soil-grown plants) and experimental treatments (phosphate starvation and AMF inoculation).

Further validation of our cluster annotations was performed using histochemical GUS staining of transcriptional reporter lines. Promoters of several genes displaying cell type-specific expression, identified using the ‘FindAllMarkers()’ function, were transcriptionally fused to GUS, and their expression patterns confirmed tissue specificity (Figure S6). Tissue type-specific expression was demonstrated for *SlWRKY72* (Solyc06g070990.3) in the epidermis; a *LIPID TRANSFER PROTEIN* (Solyc09g065430.4) in the exodermis; a *2-OXOGLUTARATE-DEPENDENT DIOXYGENASE* (Solyc06g066830.4) in the inner cortex; a transmembrane protein (Solyc12g005650.2) in the endodermis; (Solyc05g052910.4); the tomato homolog of LATERAL ROOT DEVELOPMENT 3 (*SlLRD3*; Solyc05g052910.4) in the phloem companion cells; a *SIEVE ELEMENT OCCLUSION PROTEIN* (Solyc03g111810.2) in the phloem sieve elements; and *CELLULOSE SYNTHASE 6 (SlCESA6*; Solyc02g072240.3) in the xylem.

### snRNA-seq reveals a distinctive cluster of AMF-responsive cells in tomato

To identify the cluster corresponding to root cells colonized by *R. irregularis,* the expression profiles of known early and late symbiotic genes were analyzed across the different clusters of the merged Seurat object of AMF-inoculated samples (Figures 2A and 2B). Early symbiosis-associated genes include the strigolactone (SL) biosynthesis genes ζ-carotene isomerase *DWARF 27* (*SlD27;* Solyc09g065750.3),^39^ *CAROTENOID CLEAVAGE DIOXYGENASE 7* (*SlCCD7;* Solyc01g090660.3),*^39,40^* and *CAROTENOID CLEAVAGE DIOXYGENASE 8* (*SlCCD8;* Solyc08g066650.3);^39-41^ the common symbiosis signaling genes *CALCIUM AND CALMODULIN-DEPENDENT PROTEIN KINASE* (*SlCCAMK;* Solyc01g096820.4),^42,43^ *SlCYCLOPS* (Solyc08g075760.3),^44,45^ and *SYMBIOTIC RECEPTOR-LIKE KINASE* (*SlSYMRK*; Solyc02g091590.4); and genes involved in vesicle dynamics and hyphal progression such as *EXOCYST COMPLEX COMPONENT 70 I* (*SlEXO70I;* Solyc04g077760.1),^46^ *VAPYRIN* (*SlVPY;* Solyc10g081500.1)*^47,48^* and *SlSYNTAXIN-132^49^* (Figure 2A). Late symbiotic genes include the GRAS TF-encoding *REQUIRED FOR ARBUSCULE MORPHOGENESIS 1* (*SlRAM1;* Solyc02g094340.1),*^50^* which regulates arbuscule development, and genes involved in nutrient metabolism and transport in arbusculated cells, such as *PHOSPHATE TRANSPORTER 4* and *5* (*SlPT4/5*; Solyc06g051850.2 and Solyc06g051860.3),^31,51^ *AMMONIUM TRANSPORTER* 2 (*SlAMT2*; Solyc08g067080.3), *H-ATPase 8* (*SlHA8*; Solyc08g078200.2),^52^ the ABC transporter-endocing genes *STUNTED ARBUSCULE 1* and *2* (*SlSTR1/2*; Solyc01g097430.4 and Solyc09g098410.3),^53^ *REQUIRED FOR ARBUSCULE MORPHOGENESIS 2* (*SlRAM2*; Solyc02g087500.3),^54^ which is a lipid biosynthetic gene encoding a glycerol-3-phosphate acyltransferase, and a gene encoding a MYB family TF involved in arbuscule senescence (*SlMYB1*; Solyc09g005370.1).^55^

**Figure 2.**
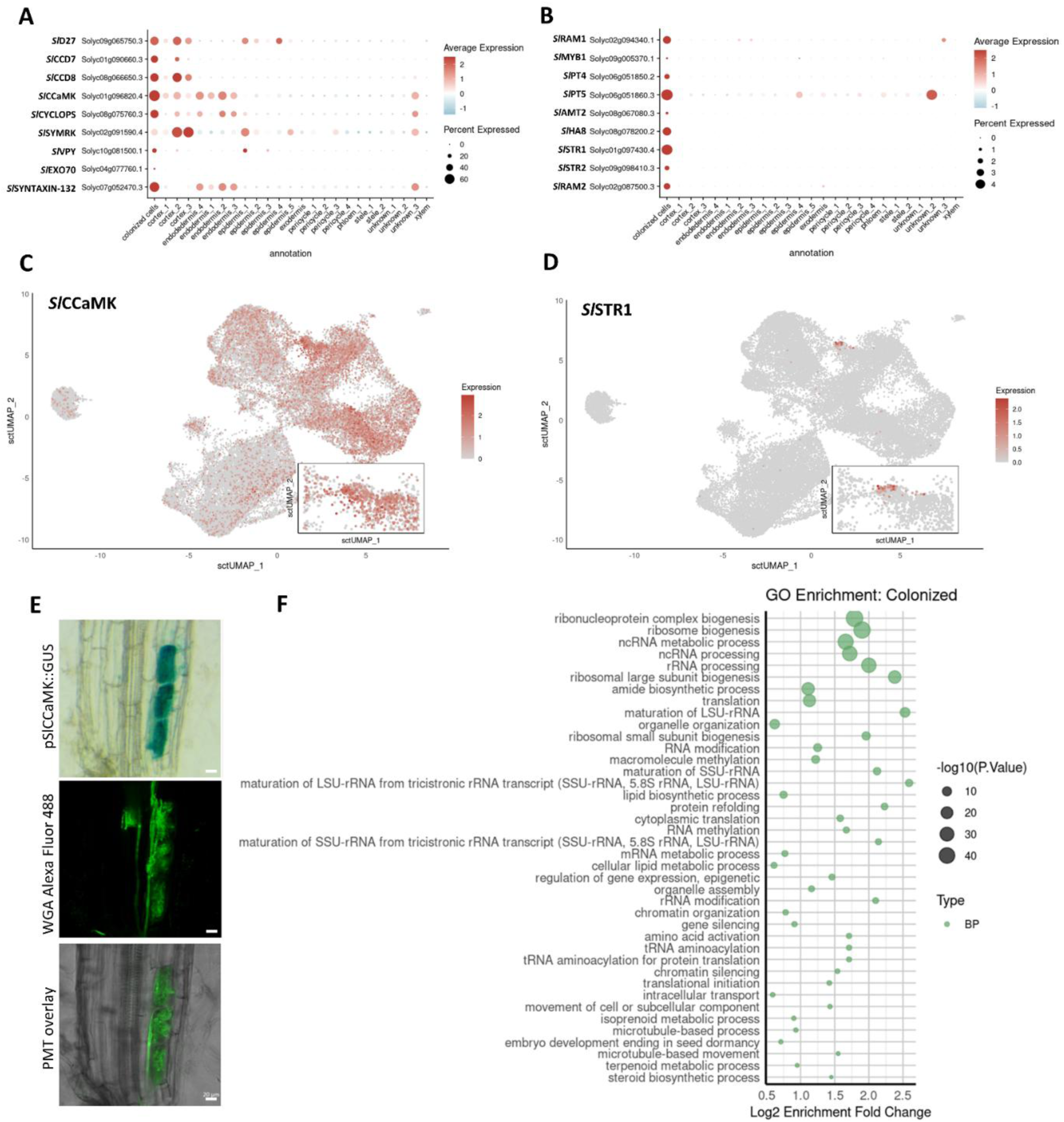
Annotation of the AMF-colonized cell cluster within the snRNA-seq dataset of AMF-colonized tomato roots. **A.** Expression profiles of well-characterized genes associated with early stages of AMF colonization, including strigolactone (SL) biosynthesis (*SlD27, SlCCD7, SlCCD8*), common symbiosis signaling (*SlCCaMK, SlCYCLOPS, SlSYMRK*), and vesicle dynamics related to hyphal progression (*SlVPY, SlEXO70, SlSYNTAXIN-132*). **B.** Expression profiles of genes associated with mature arbusculated cells, including genes involved in arbuscule development (*SlRAM1*), nutrient transport and metabolism (*SlPT4/5, SlAMT2, SlHA8, SlSTR1/2, SlRAM2*), and arbuscule senescence (*SlMYB1*). Dot size represents the percentage of nuclei expressing the gene within each cluster, while dot color indicates the average scaled expression, with red indicating higher and blue lower expression. **C-D.** UMAP projections showing expression of an early symbiotic gene (*SlCCaMK*, **C**) and a late symbiotic gene (*SlSTR1*, **D**). Color scale represents log2-normalized, corrected UMI counts. The inset zooms in on the “colonized cells” cluster. **E.** GUS staining of tomato roots from composite plants expressing *pSlCCAMK*::GUS, co-stained with WGA-Alexa Fluor 488 to visualize fungal hyphae. Scale bars = 20 µm. PMT: photomultiplier tube. **F.** GO enrichment analysis of genes significantly upregulated in colonized nuclei. Dot size represents the significance of GO term enrichment (-log10(P.value)), while the x-axis shows the log2 enrichment fold change. Only the top 40 most significantly enriched GO terms in the biological process (BP) category are shown.

Expression of early symbiotic markers, such as *SlCCaMK*, was detected in cortical, and occasionally endodermal and epidermal cells (Figures 2A and 2C). Except for *SlSYMRK*, early symbiotic genes were enriched in the cluster originally identified as cluster 16 (Figure S2), hereafter referred to as the “colonized cells” cluster, which is closely associated with cortex-related clusters (cortex_2 and cortex_3). Late symbiotic markers, such as *SlSTR1*, were almost exclusively expressed within this cluster (Figures 2B and 2D), particularly in a subregion located at its upper part (Figure 2D), supporting the interpretation that this cluster captures a developmental gradient of inner cortical cells undergoing arbuscule development. While broadly detected within the cortex, epidermis, and endodermis, *SlCCaMK* expression was significantly enriched in the “colonized cells” cluster within our snRNA-seq dataset (Figures 2A and 2C). Accordingly, transcriptional fusion of the *SlCCAMK* promoter (*pSlCCAMK) to GUS* and subsequent staining revealed predominant promoter activity in arbusculated cells, confirmed by co-staining with wheat germ agglutinin (WGA)-Alexa Fluor 488 (Figure 2E). Additionally, cluster 16 was absent from mock-inoculated samples, further implying that it encompasses root cells actively colonized by AMF (Figure S2).

Across all AMF-inoculated datasets, the “colonized cells” cluster encompassed a total of 1,154 nuclei. Identification of differentially expressed genes (DEGs) in this cluster relative to the expression in all other clusters, yielded a total of 2,843 DEGs that were significantly upregulated and 1,189 DEGs that were significantly downregulated (Figure 2F, Supplementary file S2). Upregulated DEGs included numerous known symbiosis-related genes, such as SL biosynthesis genes (e.g. *SlD27, SlCCD7*, and *Sl*CCD8), signaling components (e.g. *SlCCAMK*, *SlCYCLOPS*, and *SlSYMRK)*, hyphal progression genes (e.g. *Sl*EXO70I and *Sl*VPY), arbuscule development genes (e.g. *SlRAM1)*, and genes involved in nutrient metabolism and transport in arbusculated cells (e.g. *SlPT4/5*, *SlHA8*, *SlSTR1/2*, and *SlRAM2*).

To gain insights into the broader biological processes active in *R. irregularis*-colonized cells, a Gene Ontology (GO) enrichment analysis was performed on the upregulated DEGs of the “colonized cells” cluster (Figure 2F, Supplementary file S2). Amongst the upregulated DEGs, overrepresented GO categories included those associated with transcription and RNA-processing, and translation and ribosome biogenesis, likely reflecting the need for metabolic and structural reprogramming within arbusculated cells, necessitating increased protein biosynthesis.

Genes encoding proteins associated with lipid metabolism were equally overrepresented amongst the upregulated DEGs, including the tomato β-keto-acyl ACP synthase I-encoding gene *DISORGANIZED ARBUSCULES* (*SlDIS*; Solyc08g082620.3), potentially involved in regulating carbon supply to the fungus. *SlDIS* is a conserved AMF host gene, mutation of which results in deficient arbuscule branching in *L. japonicus* and impaired formation of lipid-containing vesicles in *R. irregularis.^56^* Other enriched lipid biosynthesis-related genes include genes encoding ACYL CARRIER PROTEINS (*SlACPs*; Solyc01g067730.3, Solyc02g038720.2, Solyc06g054510.4), a HYDROXYACYL-DEHYDRATASE (*SlHAD*; Solyc01g105060.3), an ENOYL-REDUCTASE (Solyc10g078740.2), a MALONYL-COA ACYLTRANSFERASE (*SlMCAT*; Solyc01g006980.4), and an ACETYL-COA CARBOXYLASE (*SlACCase*; Solyc09g013080.3) for malonyl-CoA production.

Additionally, GO terms related to cellular reorganization, including organelle organization and dynamics, intracellular transport and microtubule-based processes, were significantly enriched. Genes associated with these GO terms may encode proteins that facilitate hyphal progression and arbuscule development by reorienting vesicle secretion toward the symbiotic membrane. Furthermore, GO terms linked to isoprenoid and carotenoid metabolism, which are known to play a crucial role throughout AM symbiosis,^40,57^ were also significantly enriched. These GO terms included *SlD27* (Solyc09g065750.3), *SlCCD1a* (Solyc01g087250.3), *SlCCD7* (Solyc01g090660.3) and *SlCCD8* (Solyc08g066650.3).

The significantly enriched GO terms amongst genes downregulated in colonized cells were predominantly related to the plants’ perception and response to environmental stimuli, and particularly its stress response, including terms such as “response to stimulus” and “immune system process” (Supplementary file S3). This is intriguing, because these cells are closely associated with fungal hyphae, and suggests that the plants’ immune system is actively modulated to enable localized fungal colonization and symbiosis establishment.

### Subclustering reveals distinct stages of AM symbiosis progression in tomato

Expression profiling of early and late symbiotic genes (Figures 2A and 2B) indicated heterogeneity within the “colonized cells” cluster, implying the presence of nuclei from cells at different stages of AMF colonization. To explore this variation, the “colonized cells” cluster was isolated and subjected to unsupervised subclustering, revealing four distinct subpopulations (Figures 3A and 3B). To assess how these subclusters relate to the original cell populations, the subclusters were projected onto the original UMAP (Figure 3C). Based on previously outlined symbiosis-related gene expression data, we propose that subclusters 0, 1, and 3 represent progressive stages of colonization in inner cortical cells, with subcluster 3 corresponding to the most advanced stage of colonization, i.e. mature arbusculated cells (Figure 3C). In contrast, subcluster 2 is spatially distinct and aligns more closely with epidermal cells in the original UMAP. Supporting this, expression of *SlWRKY74* (Solyc06g070990.3), a putative epidermal marker in our dataset, was also detected in subcluster 2 (Figure S6).

**Figure 3.**
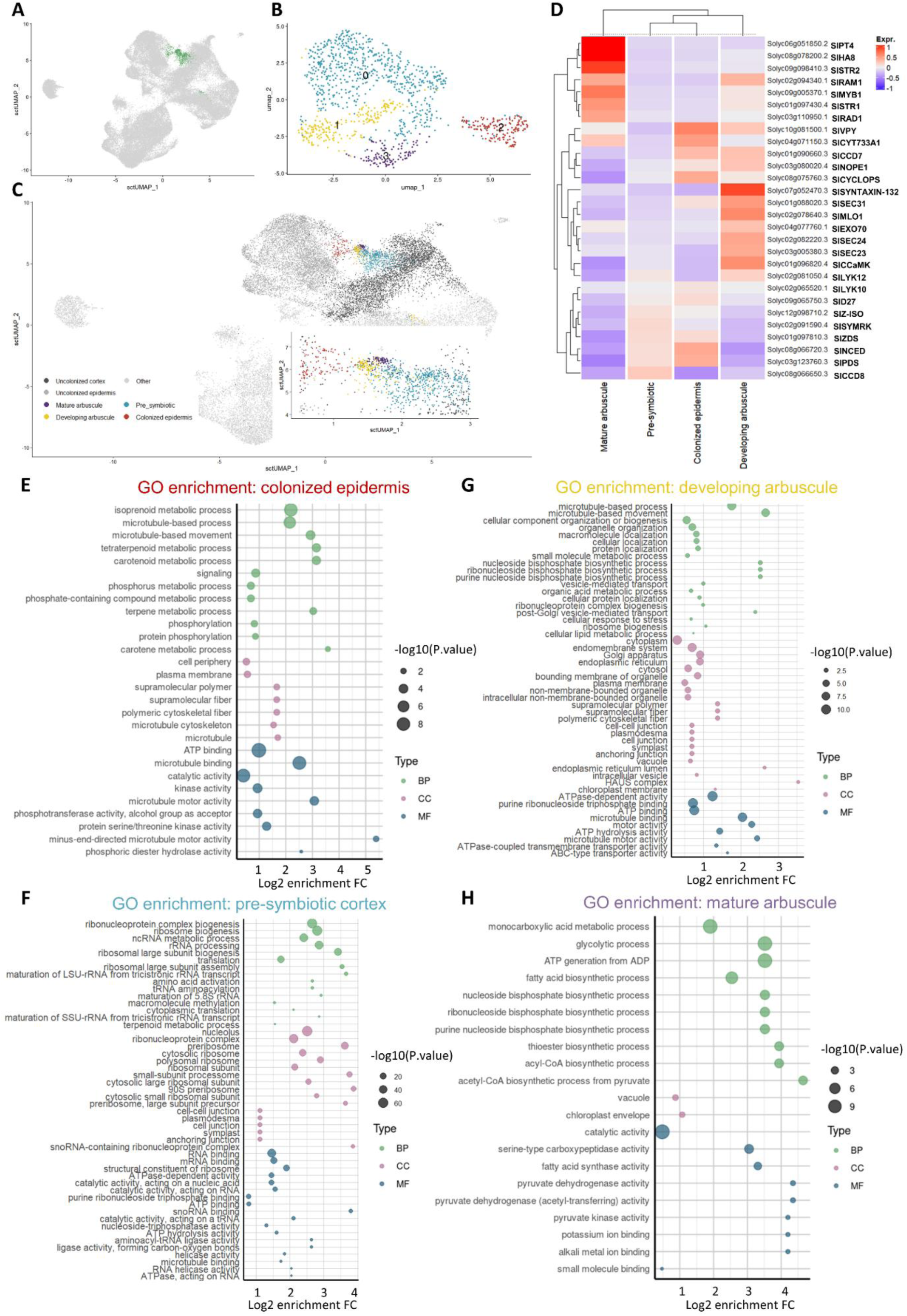
snRNA-seq unveils distinctive stages of AMF colonization in tomato roots. **A.** Depiction of the cluster representing nuclei derived from AMF-colonized cells on the snRNA-seq UMAP. Colonized cells are depicted in green. **B.** Four subclusters identified upon isolation and reclustering of the AMF-colonized cells cluster. **C.** Projection of the identified subclusters onto the original UMAP. While subcluster 0, 1, and 3 appear to correspond to progressing stages of arbuscule formation in cortical cells, subcluster 2 appears to correspond to colonized epidermal cells. The inset zooms in on the “colonized cells” cluster. **D.** Heatmap displaying the expression of established genes associated with different stages of AMF colonization, including genes involved in strigolactone (SL) biosynthesis (*SlD27, SlCCD7, SlCCD8*), common symbiosis signaling (*SlCCaMK, SlCYCLOPS, SlSYMRK*), vesicle dynamics for hyphal progression (*SlVPY, SlEXO70, SlSYNTAXIN-132*), arbuscule development (*SlRAM1*), nutrient transport and metabolism (*SlPT4/5, SlAMT2, SlHA8, SlSTR1/2*, *SlRAM2*), and arbuscule senescence (*SlMYB1*). **E.** GO enrichment analysis of the subcluster suggested to contain colonized epidermal cells (subcluster 2). **F.** GO enrichment analysis of the subcluster suggested to contain presymbiotic cortical cells (subcluster 0). **G.** GO enrichment analysis of the subcluster suggested to contain cortical cells undergoing arbuscule development (subcluster 1). **H.** GO enrichment analysis of the subcluster suggested to contain cortical cells containing mature arbuscules (subcluster 3). Only the top 50 (or less) significant enriched GO terms are displayed. Different classes of GO categories “Biological Process” (BP), “Cellular component” (CC), and “Molecular Function” (MF) are represented in green, purple and blue, respectively. The x-axis displays the log2 enrichment fold change (FC) for each GO category. Dot diameter displays the significance of the enrichment (-log10(P-Value)).

To assess functional heterogeneity within diverse subpopulations of the “colonized cells” cluster, we examined the expression of stage-specific marker genes (Figure 3D), and identified DEGs for each subcluster, followed by GO enrichment analysis for significantly upregulated genes (Figures 3E-3H, Supplementary file S4). For the colonized epidermis (subcluster 2), DEGs were determined relative to the “uncolonized epidermis”. For individual stages of inner cortical cell colonization (subcluster 0, 1, and 3), DEGs were determined relative to all other inner cortex-associated clusters of the merged AMF-inoculated dataset, including the other colonization stages and the “uncolonized cortex” (Figures 3E-3H).

In subcluster 2, a total of 647 genes were upregulated and 18 downregulated relative to the uncolonized epidermis. Enriched GO terms amongst the upregulated DEGs included signaling and phosphorylation, terpenoid and carotenoid metabolism, microtubule-based processes, and phosphorus metabolic processes (Figure 3E). Early symbiotic marker genes involved in SL biosynthesis (e.g. *SlD27* and *SlCCD7*), AMF recognition (e.g. *SlLYK10*), common symbiosis signaling pathway (CSSP) signaling (e.g. *SlSYMRK* and *SlCYCLOPS*), and vesicle trafficking for hyphal accommodation (e.g. *SlVPY*) were highly expressed within this subcluster (Figure 3D), supporting their annotation as epidermal cells preparing for, or actively undergoing, fungal entry (colonized epidermis).

Subcluster 0, appearing early in the developmental trajectory arising from the uncolonized cortex, had 1,154 up- and 450 downregulated DEGs in comparison to the remaining inner cortex (Figure 3F). GO-terms associated with upregulated DEGs included several related to ribosome biogenesis and assembly, tRNA and amino acid metabolism, RNA processing and modification, and translation and protein metabolism, suggesting active biosynthetic preparation for fungal accommodation, further supported by the concomitant enrichment of GO terms associated with microtubule-based processes. Moreover, GO terms associated with terpenoid and carotenoid metabolism were also enriched, exemplified by enhanced expression of ζ*-CAROTENE DESATURASE* (Solyc01g097810.3), *15-CIS-ζ-CAROTENE ISOMERASE* (chloroplastic; Solyc12g098710.2), *PHYTOENE DESATURASE* (Solyc03g123760.3) and *SlCCD8* (Solyc08g066650.3), specifically involved in SL biosynthesis^41^ and *9-CIS-EPOXYCAROTENOID DIOXYGENASE* (Solyc08g066720.3), involved in abscisic acid (ABA) biosynthesis^58^ (Figure 3F). Both SLs and ABA are known to promote the early stages of AMF association,^59-62^ suggesting that this subcluster represents cortical cells anticipating, or in early engagement with AMF hyphae. This is further supported by the observation that amongst the most significant and highly differentially induced genes in this cluster are genes involved in cell wall remodeling, such as *EXPANSIN 12* (Solyc05g007830.3), a *PECTINESTERASE* (Solyc06g051960.3), and a *PECTATE LYASE* (Solyc06g071840.3). Accordingly, this subcluster displayed enhanced expression of other early symbiotic markers involved in SL biosynthesis (e.g. *SlD27*, *SlCCD7*, and *SlCCD8*), symbiont recognition (e.g. *SlLYK12*), and CSSP signaling (e.g. *SlSYMRK*) (Figure 3D), reinforcing the interpretation of this subcluster as the presymbiotic cortex.

Subcluster 1, representing an intermediate colonization stage, showed 1,352 up- and 461 downregulated genes (Figure 3G). DEGs upregulated in subcluster 1 were significantly enriched in GO terms associated with subcellular reorganization and vesicle loading and trafficking, as exemplified by enhanced expression of genes encoding components of the COPII-mediated vesicle transport system *SlSEC23* (Solyc03g005380.3), *SlSEC24-LIKE PROTEINS* (Solyc02g082220.3, Solyc05g055690.3) and a *SlSEC31-LIKE PROTEIN* (Solyc01g088020.3), *MICROTUBULE ORGANIZATION 1* (*SlMOR1*; Solyc02g078640.3),^63^ several *KINESINS* (e.g. Solyc06g009780.4, Solyc08g081120.4, and Solyc11g072820.3), *KINESIN-LIKE PROTEINS* (e.g. Solyc10g083310.2, Solyc01g100120.4, Solyc07g064030.4, Solyc11g008350.2, and Solyc02g068340.3), *PHOSPHOLIPID-TRANSPORTING ATPases* (Solyc02g069430.4, Solyc02g069420.4, and Solyc05g006640.4), which maintain phospholipid asymmetry critical for membrane fusion events,^64^ and *SYNTAXIN-132* (*SlSYNTAXIN-132*; Solyc07g052470.3), a homolog of which is known to control the formation of a stable symbiotic interface in *Medicago truncatula* (Figure 3G).^49^ Moreover, GO categories related to enhanced lipid and carbohydrate metabolism were also enriched, as well as diverse categories consistent with an increased metabolic state of the cells encompassed by this cluster, including nucleotide biosynthesis and ribosome biogenesis. Accordingly, symbiotic marker genes strongly expressed in subcluster 1 include genes involved in SL biosynthesis (e.g. *SlCCD7*), symbiont recognition (e.g. *SlLYK12*), and CSSP signaling (e.g. *SlCCaMK*), but also genes involved in hyphal accommodation (e.g. *SlVPY*, *SlEXO70*, and *SlSYNTAXIN-132*), arbuscule development and functioning (e.g. *SlRAM1*, *SlRAD1*, and *SlSTR1*) and even arbuscule senescence (e.g. *SlMYB1*) (Figure 3D). Together, these data support the interpretation that this subcluster represents cells actively accommodating hyphae and forming arbuscules, requiring extensive remodeling of cellular metabolism, subcellular organization, and endomembrane trafficking.

Subcluster 3, positioned at the end of the colonization trajectory, showed 605 up- and 146 downregulated genes (Figure 3H). Enriched GO terms amongst upregulated DEGs were associated with central carbon metabolism and fatty acid biosynthesis, demonstrated by the enhanced expression of *SlRAM2* (Solyc02g087500.3), *SlSTR1* (Solyc01g097430.4), and *SlSTR2* (Solyc09g098410.3) (Figures 3D and 3H),^65,66^ and GO terms associated with nucleotide biosynthesis. Importantly, the expression of the mycorrhiza-inducible phosphate transporters (*SlPT4*; Solyc06g051850.2 and *SlPT5*; Solyc06g051860.3) and *SlHA8*, responsible for the construction of a H^+^ gradient fueling symbiotic phosphate and nitrogen (N) transport,^52^ were significantly more highly expressed in this subcluster compared to the others (Figure 3D). In contrast to earlier subclusters, expression of early symbiotic markers was markedly reduced, confirming this stage as a terminal point in the developmental progression.

Together, these findings reveal a clear functional and transcriptional stratification within the “colonized cells” cluster, representing a continuum from early symbiotic signaling and metabolic priming to active fungal accommodation and nutrient exchange. These subcluster-specific profiles provide valuable insights into the temporal dynamics of AMF colonization and offer a rich resource for prioritizing candidate genes for future functional studies.

### MINI-EX predicts TFs involved in AM symbiosis progression

To obtain a better understanding of the key TFs driving stage-specific transcriptional responses during AMF colonization in tomato roots, we applied MINI-EX, a tool for cell-type-specific gene regulatory network inference, identifying regulons in a cell-type specific manner through the integration of expression-based networks and TF-binding motif information.^67^ Candidate TFs are ranked using Borda ranking, which incorporates both network centrality measures (closeness and betweenness) and expression specificity across clusters. To identify TFs driving early symbiotic responses in the epidermis, the set of genes significantly upregulated in colonized vs. uncolonized epidermal cells was used. For cortex-associated stages of colonization (presymbiotic, developing arbuscule, and mature arbuscule), MINI-EX analysis was performed using stage-specific upregulated DEGs obtained through comparison to the rest of the cortex, as described above (Supplementary file S5).

### MINI-EX enhances our understanding of known symbiotic regulators

Application of MINI-EX on the “colonized cells” subclusters resulted in the identification of several established TFs associated with AM and/or root nodule symbiosis, providing deeper insights into their relevance and spatiotemporal activity during AM symbiosis in tomato (Figure 4). While expression of the GRAS TF-encoding tomato homolog *NODULATION SIGNALING PATHWAY 1* (*SlNSP1*; Solyc03g123400.1) was broadly detected in the uncolonized epidermis and cortex, *SlNSP1* expression was significantly increased in nuclei associated with colonized cells (Figure 4A). Our data suggests that *SlNSP1* most strongly contributes to symbiotic gene expression in epidermal cells, consistent with previous studies in *M. truncatula*, demonstrating a role for *MtNSP1* in regulating symbiotic gene expression downstream of CSSP activation and SL biosynthesis.^68^

**Figure 4.**
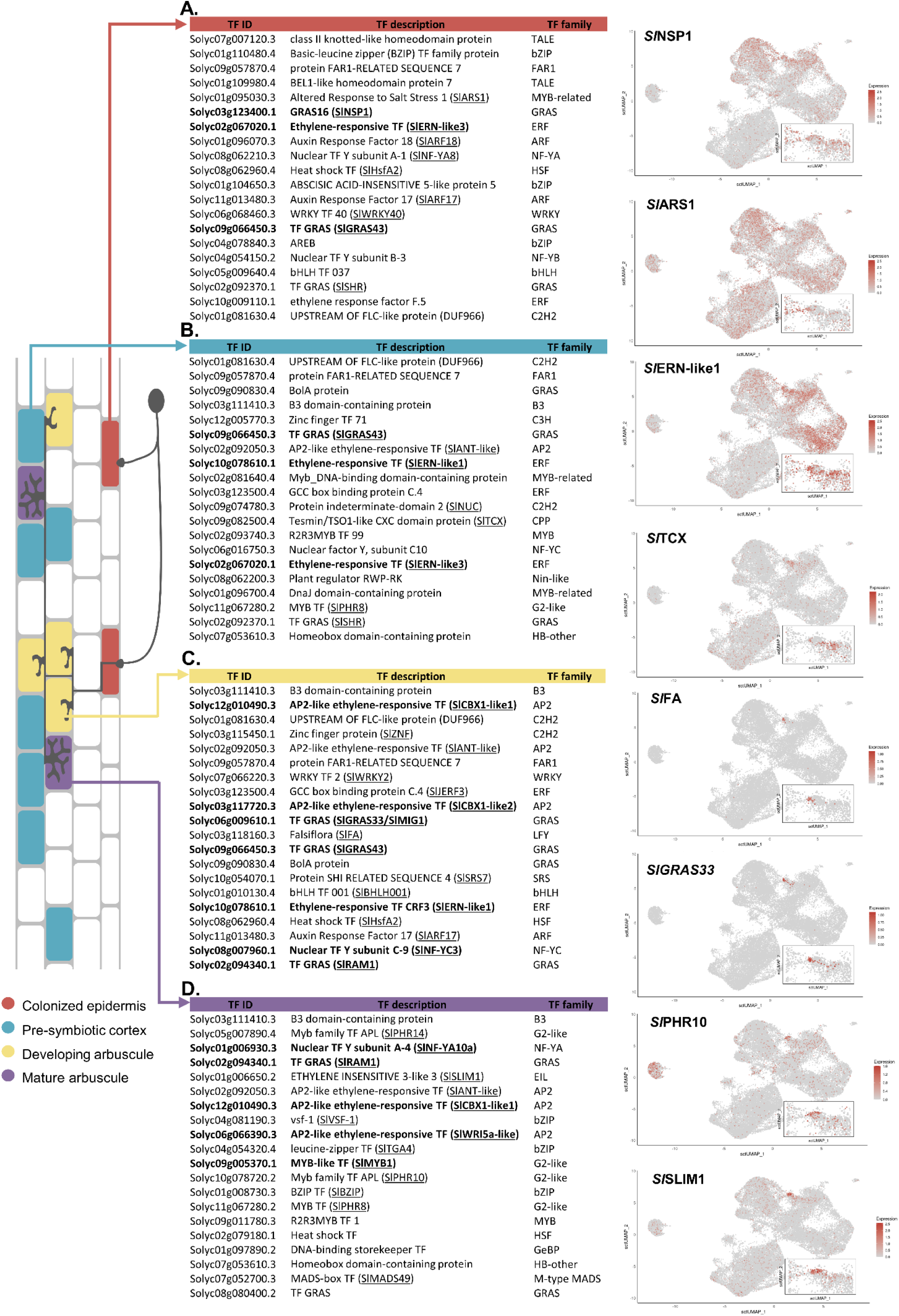
Top transcription factors (TFs) predicted by MINI-EX to drive stage-specific gene expression during AMF colonization in tomato roots. **A.** Predicted TFs active in nuclei of the colonized epidermis. **B.** Predicted TFs active in nuclei of the presymbiotic cortex. **C.** Predicted TFs active in nuclei of cortical cells undergoing arbuscule development. **D.** Predicted TFs active in nuclei of cortical cells containing mature arbuscules. All TFs are ranked according to their Borda score (ranked 1 to 20). For all predicted TFs, the TF gene ID, description and predicted TF family (PlantTFDB v5.0) is shown. For each colonization stage, UMAP expression profiles of two representative TFs are displayed on the right. The color scale represents log2-normalized corrected UMI counts. The inset zooms in on the “colonized cells” cluster. Tomato homologs of previously characterized symbiotic TFs are indicated in bold. TFs for which functional information is available (or supported through homology), albeit in a non-symbiotic context, are underlined.

We also identified *SlERN-like1* (Solyc10g078610.1) and *SlERN-like2* (Solyc10g080650.3), which are closely related to the *M. truncatula* ethylene response factor (ERF)-encoding genes *REQUIRED FOR NODULATION1/2/3* (*MtERN1/2/3*; Medtr7g085810, Medtr6g029180, Medtr8g085960) (Figure S7). In *M. truncatula* and *L. japonicus*, ERN1 facilitates infection thread formation and nodule organogenesis,^69-71^ partially redundantly regulated by MtERN2 in *M. truncatula.^69^* In contrast, the role of ERN3 during root nodule symbiosis is less documented. While functional roles for ERN1/2/3 during AM symbiosis have not yet been confirmed, ERN1/2 are induced upon AMF colonization.^70,72^ Moreover, an additional *SlERN-like3* (Solyc02g067020.1) appeared closely related to three other genes encoding AP2 TFs (Medtr6g012970, Medtr7g011630, and Medtr4g082345) (Figure S7), conserved in AM symbiosis-competent angiosperms.^73^ Both *SlERN-like1* (Solyc10g078610.1) and *SlERN-like3* (Solyc02g067020.1) were expressed throughout the epidermal and cortical clusters, with increased expression in nuclei of the “colonized cells” cluster. SlERN-like1 is predicted to primarily regulate transcriptional changes within presymbiotic cortical cells and cortical cells undergoing arbuscule development (Figures 4B and 4C), whereas SlERN-like3 appeared to primarily impact symbiotic gene expression in the epidermis.

We also detected the CCAAT-box-binding TF subunit-encoding *SlNF-YA10a* (Solyc01g006930.3), a homolog of both *MtNF-YA1* (Medtr1g056530) and *MtNF-YA2* (Medtr7g106450), previously implicated in rhizobial infection and root nodulation.^74^ In tomato, *SlNF-YA10a* was also shown to negatively regulate ascorbate biosynthesis through modulation of the D-mannose/L-galactose pathway.^75^ While broadly expressed, *SlNF-YA10a* expression was significantly enhanced in mature arbusculated cells (Figure 4C), where it is predicted to function as an important transcriptional regulator. A gene ecoding an additional CCAAT-box-binding TF subunit, *SlNF-YC3* (Solyc08g007960.1), was suggested to broadly control symbiotic gene expression in colonized epidermal cells, presymbiotic cortical cells, and especially cortical cells undergoing arbuscule development. *SlNF-YC3* is a homolog of *MtNF-YC6/MtCbf1* (Medtr2g081600) and *MtNF-YC11/MtCbf2* (Medtr2g081630), AM-conserved genes with expression patterns closely associated with fungal contact.^13,73^

Expression of the GRAS TF-encoding gene *SlGRAS43* (Solyc09g066450.3) was strongly induced in the “colonized cells” cluster, and predicted to contribute to gene expression changes in the colonized epidermis, presymbiotic cortical cells, and cells undergoing arbuscule development (Figures 4B and 4C). Although expression of *SlGRAS43* in arbusculated cells has been confirmed previously,^76^ its RNAi-mediated knock-down in tomato roots did not significantly affect mycorrhizal colonization.^76^ Another GRAS TF predicted by MINI-EX as a key regulator during arbuscule development was encoded by *Sl*GRAS33 (Solyc06g009610.1) (Figure 4C), a close homolog of *M. truncatula Mycorrhiza Induced GRAS 1* (*MtMIG1*), which facilitates radial cortical cell expansion during arbuscule development.^76,77^ Within our dataset, *SlRAM1* (Solyc02g094340.1), encoding another GRAS TF, was expressed in, and predicted to regulate transcription in, both cells undergoing arbuscule development and mature arbusculated cells, in line with previous studies highlighting RAM1’s role in arbuscule development and symbiotic nutrient exchange.^50,66,78^

Homologs of *LjCBX1,* namely *SlCBX1-like1/2* (Solyc12g010490.3 and Solyc03g117720.3) (Figure S8) were predicted to function during the later stages of AM symbiosis, with MINI-EX predicting a prominent role for both homologs in transcriptional regulation of arbuscule development (Figures 4C and 4D). Indeed, previous studies suggest that *LjCBX1* contributes to both arbuscule formation and symbiotic nutrient exchange in *L. japonicus.^79,80^* In nuclei of cells predicted to house mature arbuscules (subcluster 3), a tomato homolog of the APETALA2/ethylene-responsive factor (AP2/ERF)-domain TF *WRINKLED 5a/b/c* genes (*MtWRI5a*; Medtr8g468920, *MtWRI5b*; Medtr7g009410, *MtWRI5c*; Medtr6g011490) was also identified (*SlWRI5a*; Solyc06g066390.3) (Figure S8). In *M. truncatula*, these TFs redundantly regulate bidirectional symbiotic nutrient exchange through transcriptional activation of fatty acid biosynthesis genes (*MtPK, MtKASII, MtKAR, MtFatM*, and *MtRAM2*), lipid transfer genes (*MtSTR1* and *MtSTR2*), and nutrient transporter-encoding genes (*MtPT4* and *MtHA1*).^81^ Finally, a MYB TF with close homology to *MtMYB1* (*SlMYB1*; Solyc09g005370.1) was predicted as an important transcriptional regulator specifically within mature arbusculated cells (Figure 4D). Given the known role of MtMYB1 in arbuscule degradation,^55^ mature arbusculated cells expressing *SlMYB1* could be preparing for, or actively undergoing arbuscule senescence.

### MINI-EX predicts novel symbiotic regulators

Several additional TFs, with currently unknown roles in symbiosis, were predicted to steer gene expression across diverse symbiotic stages. This included a B3 DOMAIN-CONTAINING PROTEIN (Solyc03g111410.3), a protein of unknown function predicted as the top transcriptional regulator of cortical cells undergoing arbuscule development and those containing mature arbuscules. Another candidate was a tomato homolog of *Arabidopsis AINTEGUMENTA* (*AtANT*) (*SlANT*; Solyc02g092050.3), which regulates cell division and differentiation in leaves and floral organs, thereby controlling lateral organ size in *Arabidopsis*^82,83^(Figure 4C). A broad regulatory role in AM symbiosis across all stages – except mature arbuscules – was also predicted for a FAR1-RELATED SEQUENCE 7 PROTEIN (*SlFRS7*; Solyc09g057870.4), belonging to a group of uncharacterized proteins related to FRS transposase-derived TFs.^84^

Beyond those described above, the top 20 predicted regulators of symbiosis-induced gene expression within epidermal cells (Figure 4A) included *ALTERED RESPONSE TO SALT STRESS 1* (*SlARS1*; Solyc01g095030.3), expression of which was strongly induced in colonized epidermal cells, *AUXIN RESPONSE FACTOR 17* and 18 (*Sl*ARF17/18; Solyc11g013480.3 and Solyc01g096070.3), and *SlNF-YA8* (Solyc08g062210.3), previously implicated in N starvation responses and tomato fruit ripening.^85,86^ Additional candidates included *ABA-RESPONSIVE ELEMENT BINDING PROTEIN 1* (*SlAREB1*; Solyc04g078840.3), encoding a central player in ABA-mediated (a)biotic stress responses,^87,88^ *ABSCISIC ACID-INSENSITIVE 5-LIKE PROTEIN 5,* also termed *ABRE-BINDING FACTOR 2* (SlABF2; Solyc01g104650.3), *SlERF.F5* (Solyc10g009110.1), known to interact with *SlMYC2* in regulating jasmonic acid-induced leaf senescence,^89^ the GRAS TF-encoding *SHORT-ROOT* (*SlSHR*; Solyc02g092370.1), and *HEAT SHOCK FACTOR A2* (*SlHSFA2*; Solyc08g062960.4).

In addition to known symbiotic regulators, the top 20 predicted TFs orchestrating early AM-induced responses within the inner cortex (Figure 4B) included *JASMONATE AND ETHYLENE-RESPONSIVE FACTOR 3 SlJERF3* (Solyc03g123500.4), which also appears active during arbuscule development, and whose homologs in tobacco have been linked to carbon and reactive oxygen species metabolism.^90^ Other candidates include *SlNUC* (Solyc09g074780.3), encoding a member of the indeterminate-domain 2 family known to modulate lignin deposition in the root exodermis,^91^ *SlPHR8* (Solyc11g067280.2), encoding a predicted *PHOSPHATE STARVATION RESPONSE* TF,^92^ and a TF-encoding gene *TESMIN/TSO1-LIKE CXC DOMAIN* (*SlTCX*; Solyc09g082500.4), with high sequence similarity to *Arabidopsis TESMIN-LIKE AtTCX2* (AT4G14770) and *AtSOL1* (AT3G22760), which regulate cell division.^93,94^ A further predicted TF of interest was a RWP-RK domain-containing TF (*SlRWP-RK*; Solyc08g062200.3) of unknown function. The RWP-RK domain is frequently found in regulators of N signaling, gametogenesis, and nodule organogenesis, including NODULE INCEPTION (NIN) and NIN-like proteins (NLPs).^95^

The top 20 predicted regulators within cortical cells undergoing arbuscule development (Figure 4C) equally included a zinc finger TF of unknown function (*SlZNF*; Solyc03g115450.1), which was very specifically expressed upon arbuscule development, a WRKY family TF (*SlWRKY2*; Solyc07g066220.3), and *FALSIFLORA* (*SlFA*; Solyc03g118160.3), a known regulator of floral meristem identity, which was also highly specifically expressed in cells undergoing arbuscule development.^96^

Besides *SlNF-YA10a, SlRAM1, SlCBX1-like1, SlWRI5a*, and *SlMYB1*, MINI-EX predicted roles for several TFs with established functions in nutrient starvation responses in regulating gene expression within mature arbusculated cells (Figure 4D). A regulatory role was suggested for the PHOSPHATE STARVATION RESPONSE 10 and 14 TFs (*SlPHR10*; Solyc10g078720.2, *SlPHR14*; Solyc05g007890.4) (Figure S9). This aligns with previous studies demonstrating that PHR TFs can induce symbiotic gene expression in *M. truncatula*, *L. japonicus*, *O. sativa*, and *S. lycopersicum.^92,97-99^* Additional TFs included the ETHYLENE INSENSITIVE 3-like (EIL3) TF SULFUR LIMITATION 1 (*SlSLIM1*; Solyc01g006650.2),^100^ and TGACG MOTIF-BINDING FACTOR 4 (*SlTGA4*; Solyc04g054320.4),^101,102^ mediating sulfur and nitrate starvation responses, respectively. The suggested involvement of *SlSLIM1*, *SlTGA4*, and *SlPHR10/14* underscores the integration of multiple nutrient-sensing pathways and of the meticulous regulation of nutrient exchange in mature arbusculated cells, ensuring an optimal symbiotic outcome.

In conclusion, applying MINI-EX to the identified “colonized cells” subclusters – each representing distinct stages of AMF colonization – provides insights into the spatiotemporal regulation of known symbiotic TFs and uncovers novel candidate regulators that warrant future functional validation.

## DISCUSSION

While offering flexibility to attune AM symbiosis progression to the requirements of both symbiotic partners in a changing environment, the plastic and asynchronous nature of AM symbiosis complicates the dissection of stage-specific transcriptional regulation. The adoption of single-cell and single-nucleus RNA-seq (sc/snRNA-seq) into the plant–microbe research field enables us to tackle this complexity in an unprecedented manner.^20-22^ Here, we present a comprehensive dataset containing 65,878 high-quality nuclei, encompassing 35,357 and 30,521 nuclei of mock- and AMF-inoculated tomato roots, respectively. Arguably, aided by *pSlPT4:GFP* enrichment,^31,51^ our dataset offers a larger subset (1,154 nuclei) of AM-responsive cells than reported in previous work,^28^ providing valuable insights into the stage-specific transcriptomes associated with AM colonization of tomato roots.

Expression profiling of marker genes and GO enrichment of DEGs revealed subclusters corresponding to (1) epidermal cells responding to fungal hyphae, and a developmental trajectory of cortical cells (2) preparing for colonization, (3) actively developing arbuscules, and (4) containing mature, functional arbuscules. Many known symbiotic genes were found among the upregulated DEGs in these subclusters, enhancing our understanding of their spatiotemporal regulation during AM symbiosis in tomato. In addition, this dataset also represents a valuable resource for the selection of novel candidate genes for further functional characterization. Furthermore, MINI-EX predicted stage-specific regulons orchestrating symbiotic gene expression in mycorrhizal tomato roots. While multiple novel candidate regulators of AMF colonization progression were proposed, our analysis also recovered a substantial number of previously described and characterized symbiotic TFs, or close homologs thereof, providing a solid proof-of-concept for our approach (Table S2).

### Regulation of strigolactone biosynthesis and the common symbiosis signaling pathway during early symbiosis

Our snRNA-seq dataset revealed two distinct subclusters representing early stages of AM colonization: epidermal cells responding to fungal signals and presymbiotic cortical cells, which represent a developmental transition from the inner cortex towards the colonized cortex. Both stages show strong expression of a subset of CSSP genes,^1^ alongside genes involved in carotenoid biosynthesis. Notably, the latter includes key enzymes for ABA and SL production, hormonal regulators known to facilitate initial symbiotic signaling and fungal recruitment.^59-62^

This finding extends the paradigm that “early” symbiotic responses are confined to the epidermis, highlighting their continued activity within the root cortex, where they may mediate fungal recruitment and preparation of cortical cells for fungal invasion. Indeed, inner cortical cells are known to induce prepenetration apparatus formation and cell-cycle reactivation prior to fungal colonization.^103-106^ Moreover, our data suggest that expression of CSSP-related and carotenoid biosynthesis genes persists in cortical cells until mature arbuscule formation is completed. This is in accordance with previous studies suggesting a role for CSSP-involved genes (e.g. *CCaMK*, *CYCLOPS*, and *VPY*) in arbuscule development.^47,78,107^

The majority of MINI-EX predicted TFs associated with early symbiotic responses in the epidermis and presymbiotic cortex remain to be functionally characterized. However, several of the predicted TFs were previously described for their role in symbiosis establishment. For example, *SlNSP1* is known to activate SL biosynthesis directly downstream of CSSP signaling in *M. truncatula*, *L. japonicus*, and rice.^68,108^ Similarly, *SlERN-like* TFs – previously shown to induce early symbiotic genes during nodulation in both the epidermis and cortex^69^ – may have AM-related functions that warrant further investigation.

### Regulation of cell division, growth, and membrane trafficking during arbuscule development

One “colonized cells” subcluster, situated midway along the AM-responsive inner cortical cell developmental trajectory, displayed the upregulation of many genes involved in intracellular rearrangement, endomembrane system reorganization, and microtubule-associated processes. Because arbuscule development is known to require substantial cellular restructuring to support the formation of the peri-arbuscular membrane (PAM) and facilitate fungal accommodation,^7,46,109^ this subcluster likely represents cells actively undergoing arbuscule development. Accordingly, MINI-EX predicted the involvement of tomato homologs of *Lj*CBX1 and *Mt*MIG1 in orchestrating transcriptional changes in these cells. Previous studies have suggested that *Lj*CBX1 contributes to arbuscule development by transcriptional regulation of a PAM-associated *Lj*KIN3-*Lj*AMK8/24 complex, besides its established role in mediating the symbiotic nutrient exchange.^79,80,110^ A role in arbuscule development was suggested for *Mt*MIG1, which is specifically expressed within arbusculated cells and recruits *Mt*DELLA1 to induce the transcription of genes involved in radial cell expansion, thought to facilitate arbuscule accommodation.^77^ Accordingly, knockdown of *MtMIG1* resulted in malformed arbuscules.^77^ Our data support a conserved involvement, and spatiotemporal expression pattern of *SlCBX-like1/2* and *SlMIG1* during arbuscule development in tomato.

Previous studies have shown that inner cortical cells reactivate the cell cycle to enable arbuscule accommodation, with arbuscule formation correlating with ectopic cell divisions and endocycling, often resulting in a higher nuclear ploidy of arbusculated cells.^103,105,106^ This reactivation likely supports arbuscule development by facilitating centripetal exocytosis, resembling cell plate formation,^111^ and by increasing cell size and biosynthetic capacity through endoreduplication.^106^ While the molecular regulators remain unclear, inspiration can be drawn from the evolutionarily related root nodule symbiosis in legumes, which also requires both cell division and endoreduplication for microbe accommodation.^112^ For instance, the *MtSHR–MtSCR* module promotes cortical cell divisions in *M. truncatula* nodules.^113^ Although its role in AM symbiosis is unconfirmed, our data implicate tomato SlSHR in transcriptional reprogramming of colonized epidermal and cortical cells. Like AtSHR and MtSHR, SlSHR drives periclinal divisions in tomato roots,^36^ making it a strong candidate for functional studies in AM symbiosis.

The predicted roles of tomato *AINTEGUMENTA-like* (*SlANT-like*) and *FALSIFLORA* (*SlFA*), close homologs of *Arabidopsis AtANT* and *LEAFY* (*AtLFY*), in arbuscule development are also intriguing. In floral meristems, AtLFY/SlFA is known to govern floral identity, while AtANT promotes organ growth via cell proliferation and expansion.^83,96,114,115^ Moreover, the here reported *SlANT-like* gene was also previously reported to be expressed in root tips.^116^ Interestingly, AtANT induces transcription of *AtLFY* in *Arabidopsis*, spatially linking the activity of the two TFs in a developmental context.^117^ Although shoot meristem genes have not been linked to AM symbiosis, several have been co-opted in root nodule development, with *M. truncatula LIGHT SENSITIVE HYPOCOTYL 1/2* (*MtLSH1/2*) and *NODULE ROOT 1/2* (*MtNOOT1/2*), orthologous to the *Arabidopsis* genes *BLADE-ON-PETIOLE 1/2* (*AtBOP1/2*), being required for root nodule meristem identity.^118-120^ Moreover, *MtLSH1/2* drives cell division to accommodate rhizobia, potentially through promotion of endoreduplication and maintenance of cells in an undifferentiated state.^119^ Given the established role of cell cycle reactivation and endoreduplication in arbuscule development, it is tempting to speculate that the floral meristem genes *SlANT-like* and *SlFA* may fulfill similar roles during arbuscule development in tomato.

### Integration of nutritional ques to finetune AM colonization and arbuscule development

At the end of the AM-responsive inner cortical cell developmental trajectory, we identified a subcluster representing nuclei of functional, mature arbuscules, supported by the specific expression of symbiotic nutrient transporters (e.g. *SlPT4* and *SlSTR1/2*) within this subcluster.^31,53^ Accordingly, GO enrichment analysis suggested that these cells display a heightened metabolic state, with for example genes involved in C-metabolism being overrepresented.

Consistent with their role in nutrient exchange, transcriptional regulation within arbusculated cells appears to be dominated by TFs involved in nutrient sensing and starvation responses. Our analysis supports roles for tomato homologs of LjCBX1 and MtWRI5a in regulating arbuscule functioning, likely through induction of nutrient transporters (e.g. SlPT4 and SlSTR1/2) and lipid metabolism-related genes.^80,81,110^ Additionally, MINI-EX analysis implicated known nutrient starvation-response TFs in AM symbiosis in tomato, particularly members of the MYB PHR1-like TF family. SlPHR8, SlPHR10, and SlPHR14 were predicted to drive gene expression changes within colonized cells, especially those containing arbuscules. These findings align with previous studies in *M. truncatula, L. japonicus, O. sativa*, and tomato,^92,97-99^ although the PHRs identified here differ from an earlier tomato study that highlighted *SlPHR1, SlPHR4, SlPHR11*, and *SlPHR12.^92^* Albeit being expressed in nuclei corresponding to colonized cells, expression of *SlPHR1/4/11* and *12* was not enhanced within these nuclei in our study. *SlPHR1* (Solyc05g055940.3) appeared generally expressed within mycorrhized roots, while the expression of *SlPHR12* (Solyc10g083340.3) and *SlPHR4* (Solyc02g067160.4) appeared to be restricted to the phloem, or more generally to the stele, respectively. In contrast, *SlPHR11* (Solyc10g080460.2) was predominantly expressed in epidermal cell clusters. This suggests functional redundancy and stage-specific roles among PHRs, warranting further investigation into their spatiotemporal dynamics.

In addition, the TF SlTGA4 was also upregulated in mature arbusculated cells and predicted by MINI-EX to be a key regulator of symbiotic gene expression. Its *Arabidopsis* homolog*, AtTGA4,* is drastically upregulated upon N deficiency, and activates genes involved in the N starvation response, including nitrate transporter genes (*AtNRT2.1* and *AtNRT2.2*).^101,102^ Furthermore, SlEIL3/SlSLIM1 was also predicted to be an important regulator of symbiotic gene expression within mature arbusculated cells.^100,121,122^ In both *Arabidopsis* and tomato, SLIM1 has been put forward as a central regulator of the sulfate (S) starvation response, with *SlSLIM1* being strongly upregulated upon S deficiency in tomato.^100^ *SLIM1* induces the expression of genes involved in S acquisition, such as sulfate transporters (*SULTRs*), as well as the degradation of glucosinolates.^121,123^ As AMF aid in the host plant’s S supply, SlSLIM1 could be implicated in the induction of *SULTR*s, and the modulation of S metabolism.^16,124,125^ However, the potential role of *SLIM1* in the context of symbiosis could be more complex, with literature suggesting that the S starvation response is tightly linked with sugar metabolism and phosphate nutrition.^100,122,126^

### Remarks and future perspectives

This dataset enables us to explore the *R. irregularis*–tomato interaction at high resolution, distinguishing multiple stages of AM colonization and revealing key genes and TFs orchestrating these processes. However, the proportion of nuclei originating from mature arbusculated cells remained relatively low. A similar limitation was also evident in the previously published snRNA-seq dataset from *M. truncatula* roots colonized by AMF, where only a small number of *MtPT4* and *MtSTR1/2*-expressing cells were detected.^28^ Notably, as in our study, their data also lacked a clearly resolved cluster corresponding to arbuscules undergoing degradation, which are expected to represent a distinct transcriptional state.^55^

While our current approach has the advantage of offering a global perspective of how different tomato root cell types respond to AMF colonization, the resolution of stage-specific transcriptional signatures in AM symbiosis could be further improved by implementing fluorescence-activated sorting of nuclei associated with active colonization prior to sequencing.

This approach would enable selective enrichment of nuclei from AM-responsive cells, provided that nuclear fluorescence is driven by promoters with strong and stage-specific activity across different stages of arbuscule development. Such targeted approaches could allow for more comprehensive profiling of the symbiotic root cortex, potentially resolving colonization stages that are notoriously rare – such as arbuscules undergoing degradation – or identifying yet uncharacterized stages.

Altogether, our dataset and MINI-EX analysis provide a detailed transcriptional landscape of the *R. irregularis*–tomato interaction, illuminating stage-specific regulatory programs and the transcriptional regulators that guide them. This work lays a robust foundation for probing how developmental and physiological cues are integrated during AM symbiosis and highlights intriguing candidate regulators for future functional validation.

## MATERIAL AND METHODS

### Composite plant generation and growth conditions

Tomato (*S. lycopersicum*) cv. Moneymaker seeds were surface-sterilized in 4% bleach for 10 min, washed with sterile demineralized water and pregerminated on moisturized Whatman filter papers for five days in the dark, at 24°C.

Composite plants were generated using *Agrobacterium rhizogenes* (strain K599)-mediated hairy root transformation. To this end, a third of the root of 5-day-old tomato seedlings was cut off diagonally, and the freshly cut surface was coated with a culture of transformed *A. rhizogenes* K599 grown on agar plates of Yeast Extract Beef (YEB) medium supplemented with 100 mg/L spectinomycin (Sp) and 300 mg/L streptomycin (St). Transformed seedlings were grown vertically on square Petri dishes with 1/2 Murashige and Skoog (1/2 MS, without sucrose) for 4 weeks at 24°C under long-day conditions (16h light, 8h dark). Plantlets were screened weekly for transformant roots using a *pRolD*:*mRuby-NLS* screening module, with weekly excision of non-transformed roots and transfer to fresh 1/2 MS plates.

At four weeks post transformation, composite plants were individually transferred to 50-ml falcons containing a sand-vermiculite mixture (1:1). AMF-inoculated plants were supplemented with *R. irregularis* DAOM197198 (approx. 250 spores per plant, Symplanta, Germany). The composite tomato plants were grown under greenhouse conditions (22 h light, 6 h dark) at 24°C until 6 wpi. The plants were watered twice weekly with modified Hoagland solution, containing 20 µM KH_2_PO_4_ until harvesting.^127^

### Cloning of gene constructs

To generate the transcriptional GUS/GFP-fusion constructs, the promoter sequences of *pSlPT4*, *pSlCCaMK*, and cluster marker genes (*Solyc06g070990.3, Solyc09g065430.4, Solyc06g066830.4, Solyc12g005650.2, Solyc05g052910.4, Solyc03g111810.2, Solyc02g072240.3*) were domesticated from tomato gDNA using promoter region-specific primer pairs (Table S3), flanked by complementary overhangs for the Golden Gate pGGA entry module. For each of the included genes, we aimed to amplify the 3-kb promoter, trimming if necessary due to the vicinity of other genes. pGGA entry vectors were predigested using BsaI, and Gibson assembly was used to insert the amplified promoter sequences.^128^ For pGGA modules containing the promoter of a cluster marker gene, Golden Gate assembly was performed with other entry modules: pGGB-GFP, pGGC-GUS, pGGD-linker, pGGE-35ST and pGGF-*pRolD:mRuby-NLS*, in the pGGP-AG destination vector.^129,130^ In the case of transcriptional *GUS* fusions for pGGA-*pSlPT4* and pGGA-*pSlCCaMK*, pGGB-*GFP* was replaced with pGGB-linker to allow compatibility with WGA-Alexa Fluor 488 staining of fungal hyphae. For the *pSlPT4:GFP* construct used for enrichment of actively colonized roots prior to snRNA-seq, the entry vectors pGGB-linker and pGGC-GFP were used.

### GUS and WGA-Alexa Fluor 488 staining

For GUS staining, roots of composite plants expressing *promoter:GUS* constructs (*pSlCCaMK:GUS*, *pSlPT4:GUS* and transcriptional fusions of cell-type-specific promoters) were harvested at 6 wpi. Roots were washed in NT buffer (50 mM NaCl, 100 mM Tris, pH 7), and subsequently vacuum infiltrated in GUS buffer (2 mM K₃[Fe(CN)₆] and 2.5 mM X-gluc in NT buffer) for 15 min, followed by a 1-4 h incubation (depending on the construct) in GUS buffer at 37°C in the dark. GUS-stained roots were incubated in 50% EtOH until imaging or further processing. After staining, roots expressing transcriptional fusions of root cell type markers were embedded in 5% agarose solution and sectioned using the Leica VT1200 vibrating blade microtome into 150-μm thick sections, which were mounted on glass slides and imaged using the Olympus BX51 light microscope.

In case of *pSlCCaMK:GUS* and *pSlPT4:GUS*, co-staining with WGA-Alexa Fluor 488 was performed for visualization of fungal structures. To this end, GUS-stained roots were rinsed with phosphate-buffered saline (PBS) and incubated in 0.1M HCl for 2h at RT. Subsequently, roots were washed with PBS and incubated for 2 h in 10 µg/ml Wheat Germ Agglutinin (WGA)-Alexa Fluor 488 (Invitrogen) in PBS at RT. Roots were washed in PBS and imaging was performed using the Zeiss LSM710 confocal microscope.

### Nuclei extraction from tomato roots

AMF- and mock-inoculated composite tomato cv. Moneymaker plants with roots coexpressing *pRolD:mCherry* and *pSlPT4:GFP* were harvested for nuclei extraction at 6 wpi. Prior to harvesting, successful transformation of mock- and AMF-inoculated roots was validated by screening for *pRolD:mCherry* expression using the Leica M165 FC fluorescence bino. In addition, abundantly colonized AMF-inoculated roots were enriched through screening for roots with abundant cortical cells displaying *pSlPT4:GFP* expression. Screening and harvesting were performed on ice. Per repeat, root sections of a total of 10 to 12 plants were pooled, in order to obtain a total of approximately 200 mg of root material. Harvested root material was placed in the lid of a glass Petri dish and chopped intensively in 1 ml of nuclei extraction buffer (NEB; 10 mM MES-KOH pH 7, 10 mM NaCl, 10 mM KCl, 2 mM EDTA, 250 mM mannitol, 0.1 mM spermine, 0.5 mM spermidine, 0.4 units/µl RNase inhibitor - 50:50 mix of Invitrogen SUPERase-In^TM^ and Promega RNAsin Plus) using a razor blade for 1 min on ice, followed by 1 min of incubation. The chopped root material was strained through a 40-µm strainer (pluriSelect), together with 1 additional ml of NEB to collect remaining debris on the Petri dish. Nuclei were stained using 40 µl of propidium iodide, and sorted using the BD Biosciences FACSDiscover™ S8 Cell Sorter.

### Single-cell library preparation and sequencing

For each sample, between 77,000 and 300,000 nuclei were sorted using the BD Biosciences FACSDiscover™ S8 Cell Sorter (Supplementary file S1). Afterwards, nuclei were centrifuged at 4°C for 10 min at 700 g using a swinging bucket rotor, resuspended in nuclei buffer and around 30,000 nuclei were loaded on a Chromium X Single Cell Instrument (10x Genomics). In the first replicate, the Chromium Next GEM Single Cell 3ʹ Reagent Kit v3.1 (10x Genomics) was used, while in the other two replicates the Chromium GEM-X Single Cell 3’ Kit v4 was used, both according to the manufacturer’s instructions. Sequencing libraries were loaded on an Illumina NovaSeq6000 flow cell and sequenced following recommendations of 10x Genomics at the VIB Nucleomics Core (VIB, Leuven).

### Raw data processing

The FASTQ files obtained after demultiplexing were used as input for “cellranger count” (version 8.0.1, 10x Genomics) to map the reads to the *S. lycopersicum* reference genome, found on PLAZA.^131^ Initial cell numbers and reads per cell are reported in Supplementary file S1. Preprocessing of the data was done by the scater (v1.20.1) R package according to the workflow proposed by the Marioni lab.^132^ Outlier cells were defined as having fewer than 500 expressed genes or 600 UMIs, or having more than 15% mitochondrial or 15% chloroplast transcripts. Further data analysis was performed with Seurat v4. Samples were merged, normalized, and scaled using the SCTransform function with the top 3000 variable features. Principal components were calculated and batch effects were corrected using Harmony.^32^ The UMAP was generated using the top 25 harmony dimensions and a resolution of 0.8 was used for the FindClusters function. For subclustering of the “colonized cells” cluster, a subset consisting of cluster 16 originating from all AMF-colonized samples was created, normalized, and scaled using the SCTransform function with the top 3000 variable features, and principal components were calculated. The UMAP was generated using the top 20 harmony dimensions and a resolution of 0.2 was used for the FindClusters function.

### MINI-EX

The prediction of TFs driving cluster-specific gene expression was performed using MINI-EX v3.0, a tool implemented as a Nextflow v24.10 pipeline and distributed via a Singularity v4.3.0 container,^67^ with added support for analysis of *S. lycopersicum* provided in the ‘data_sly’ folder. A list with putative S. lycopersicum TFs—i.e. proteins predicted to contain a DNA-binding motif—was obtained from the Plant Transcription Factor Database (PlantTFDB v5.0). The pipeline was executed with default parameters, with the following modifications: motif enrichment analysis was disabled, and a standard ranking procedure was used without prioritization based on predefined terms of interest. Additionally, the ‘expressionFilter’ parameter was set to 2 to increase sensitivity and allow detection of lowly expressed transcription factors.

### Phylogenetic tree construction

For phylogenetic tree construction, amino acid sequences of the AP2/ERF domain homologous gene families HOM05D001032 (ERN-like) and HOM05D000138 (WRI5/CBX1-like), as well as the MYB domain homologous gene family HOM05D000184 (PHR1-like), were retrieved from PLAZA 5.0. Sequences were aligned using MUSCLE embedded in MEGA12, and phylogenetic trees were inferred using the Maximum Likelihood (ML) method implemented in MEGA12. The JTT+Freq model was used for HOM05D001032 and HOM05D000184, whereas the JTT model was applied for HOM05D000138.

For each dataset, the tree with the highest log-likelihood is presented. Adaptive bootstrapping was performed, and bootstrap support values were estimated for each tree. Initial heuristic trees were selected based on log-likelihood comparisons between Neighbor-Joining (NJ) and Maximum Parsimony (MP) methods. Rate variation among sites was modeled with a discrete Gamma distribution (+G), and the proportion of invariant sites (+I) was estimated for each dataset.

## ACKNOWLEDGMENTS

We thank Dr. Juan A. López-Ráez for the inspiring discussions in the framework of the ROOT-BENEFIT COST (European Cooperation in Science and Technology) action (CA22142) and Dr. Annick Bleys for critically reading and finalizing the manuscript. We are also thankful for the VIB Single Cell Core, VIB Flow Core Ghent and VIB Nucleomics Core for support and access to the instrument park (vib.be/technologies). This research was supported by the Concerted Research Actions fund of Ghent University (BOF18-GOA-013) to S.G, and by Research Foundation Flanders (FWO-Vlaanderen) through predoctoral Basic Strategic Research fellowships to N.S. (1S14621N) and T.L. (1S09624N), a postdoctoral fellowship to J.V.D. (1279524N), and project G045921N awarded to P.V.D. T.E. was supported by the European Research Council (ERC StG TORPEDO; 714055 and ERC CoG PIPELINES; 101043257 to B.D.R).

## AUTHOR CONTRIBUTIONS

N.S., T.L., J.V.D. and S.G. conceptualized the research. N.S., T.L., A.D.K. and T.E. performed the experimental work. N.S. and T.E. analyzed the data. N.S. wrote the manuscript. All authors revised the manuscript. S.G., J.V.D., P.V.D. and B.D.R., supervised the research. S.G., J.V.D. and P.V.D. supervised and edited the writing.

## SUPPLEMENTAL INFORMATION

Supplemental information can be found online.

## DATA AVAILABILITY

The snRNA-seq data generated in this study will be publicly released via NCBI GEO upon acceptance of the manuscript for peer-reviewed publication.

