## Supplementary figures and tables for "Decoding stage-specific symbiotic programs in the *Rhizophagus irregularis*–tomato interaction using single-nucleus transcriptomics"

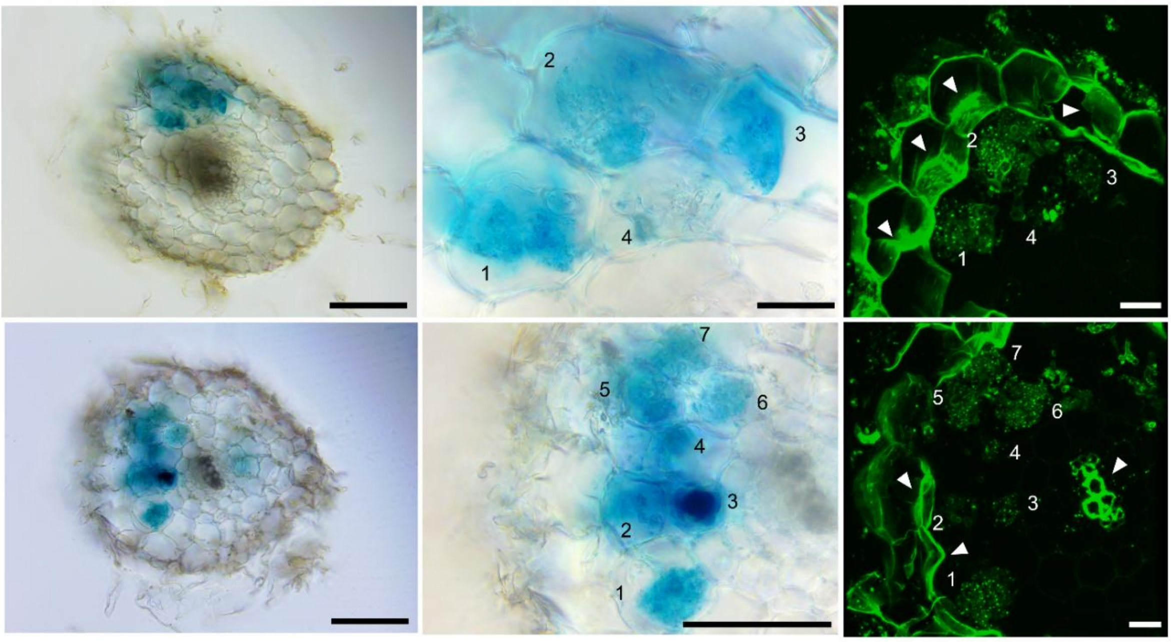


**Figure S1. The *pSlPT4* promoter activates transcription in arbusculated tomato root cells.** Composite plants with roots expressing *pSlPT4*:*GUS* were generated using *Agrobacterium*-mediated hairy root transformation. Composite plants were inoculated with *Rhizophagus irregularis* spores and imaged at 6 weeks post inoculation (wpi). *pSlPT4:GUS*-stained roots were imaged using the Olympus BX51 microscope. WGA-Alexa Fluor 488 was used for imaging of fungal structures, and confocal images were taken using the Zeiss LSM710. *pSlPT4:GUS*-stained cells and corresponding arbuscules in the WGA-stained image are labeled with the same number. For the confocal images, visualization of arbuscules in multiple planes was achieved using a Z-stack, which was assembled into a 3D project using ImageJ. Scalebars of GUS-stained roots are 100 μm, except for the top right image, where the scale bar is 20 μm. Scalebars of WGA-Alexa Fluor 488-stained roots are 20 μm. Regions of high autofluorescence, corresponding to exodermal and xylem cells, are indicated with white arrowheads. This autofluorescence is likely due to suberin and lignin accumulation in secondary cell walls of these cell types.

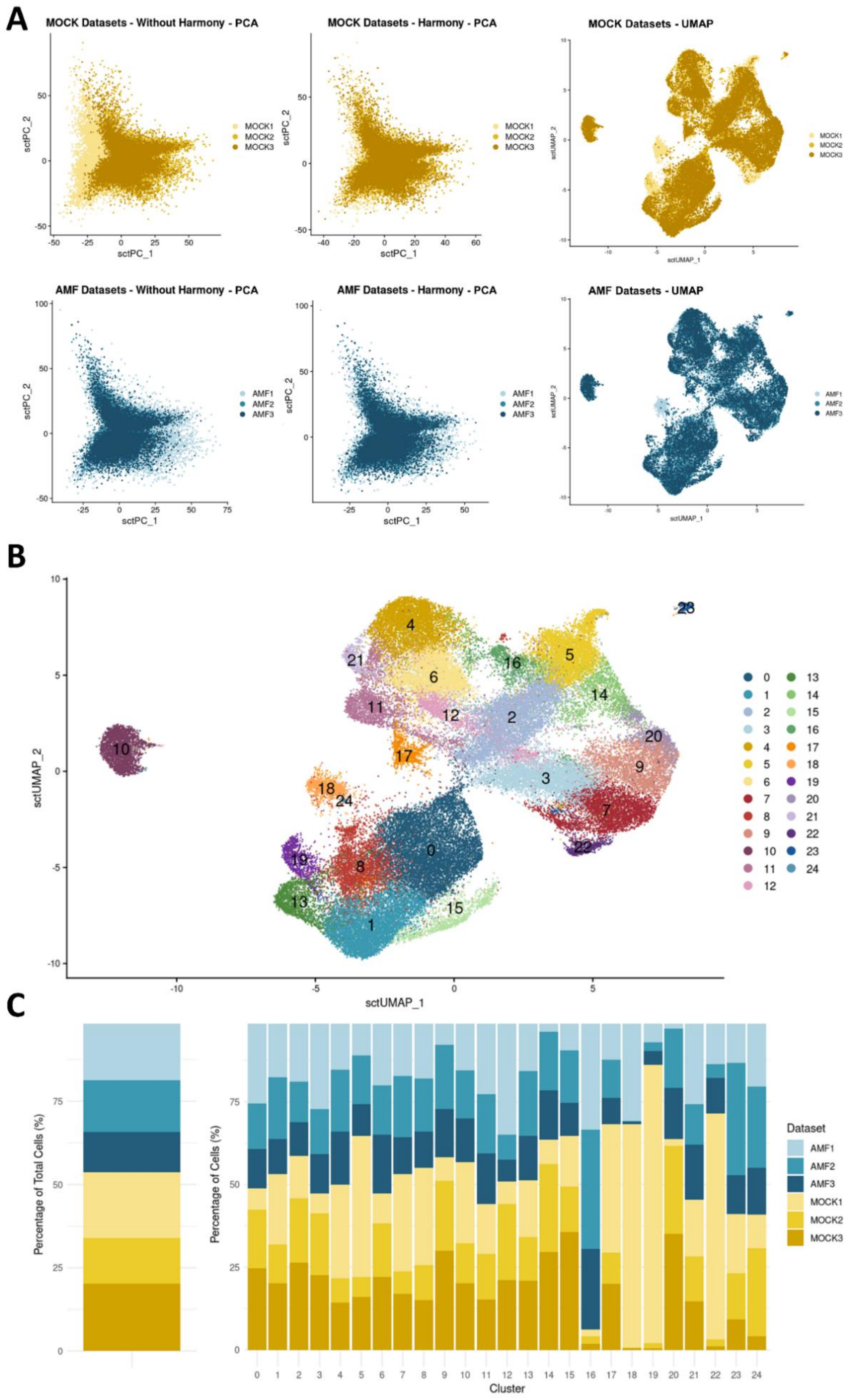


**Figure S2. Contribution of individual snRNA-seq datasets to the final snRNA-seq tomato–*R. irregularis* dataset.** **A.** Overlap of principal component analysis (PCA) plots of individual MOCK and AMF datasets prior to and after Harmony batch correction, and in the final UMAP plot. **B.** UMAP of the merged snRNA-seq dataset of MOCK and AMF-inoculated roots with 24 individual clusters identified using unsupervised clustering. **C.** Contribution of individual datasets to the final snRNA-seq dataset and its individual clusters.

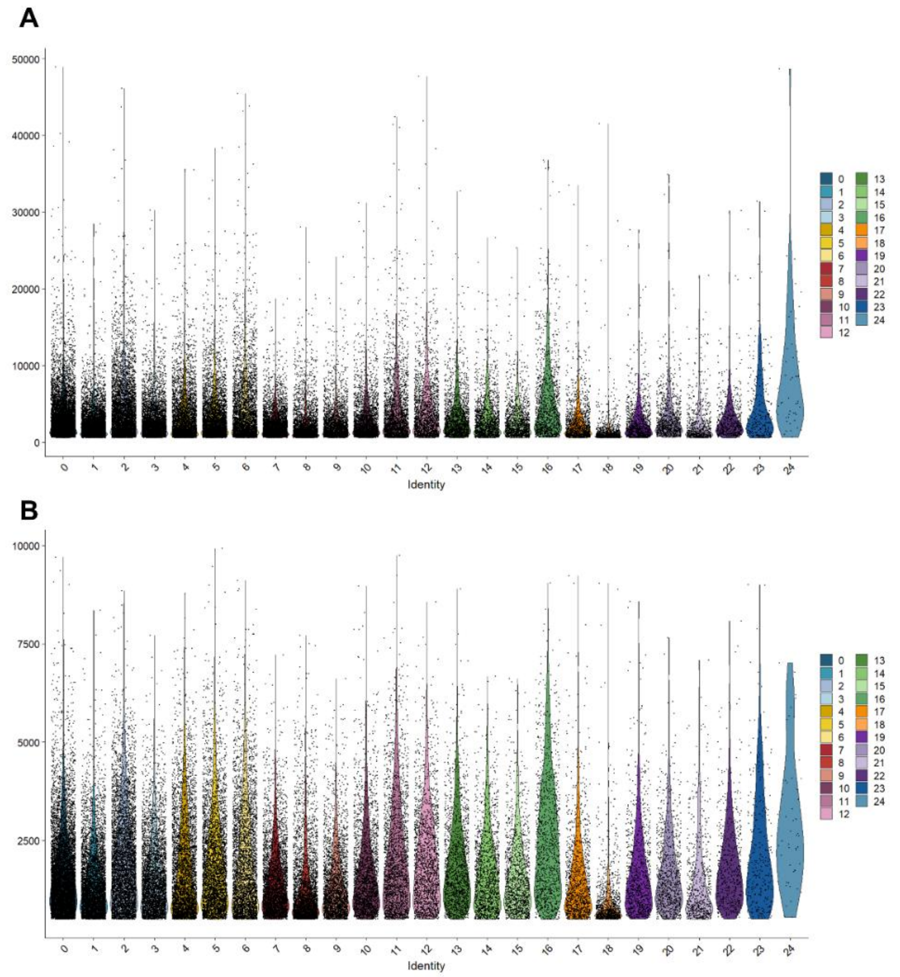


**Figure S3. Violin plots representing the total number of transcripts (A) and genes (B) detected per nucleus for each cluster of the merged snRNA-seq dataset of MOCK and AMF-inoculated tomato roots.**

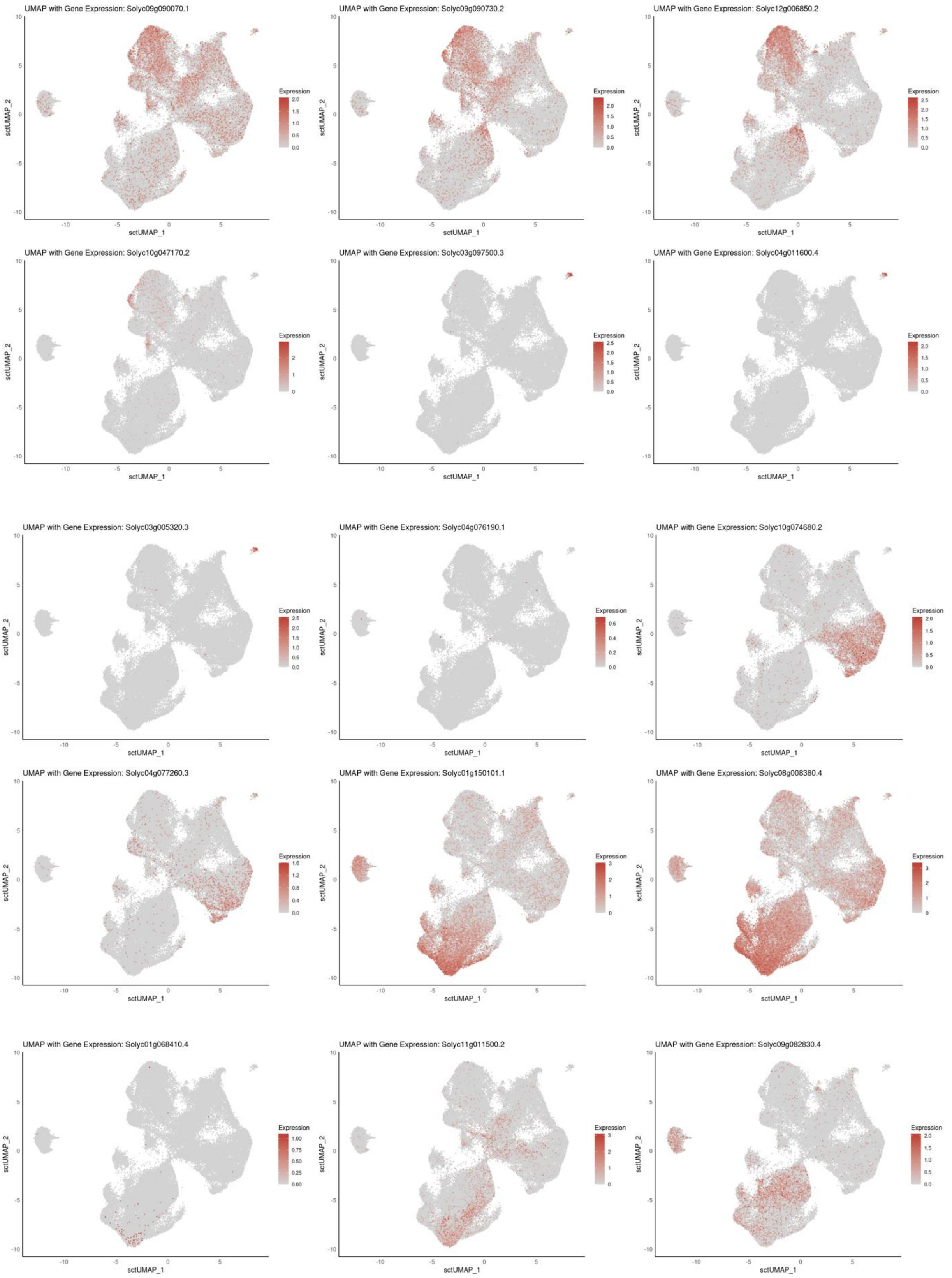


**Figure S4. The expression of marker genes of individual clusters displayed on a UMAP of the snRNA-seq tomato–*R. irregularis* dataset**. Dots represent cells colored by normalized mRNA counts (expression). The utilized marker genes are summarized in Table S1.

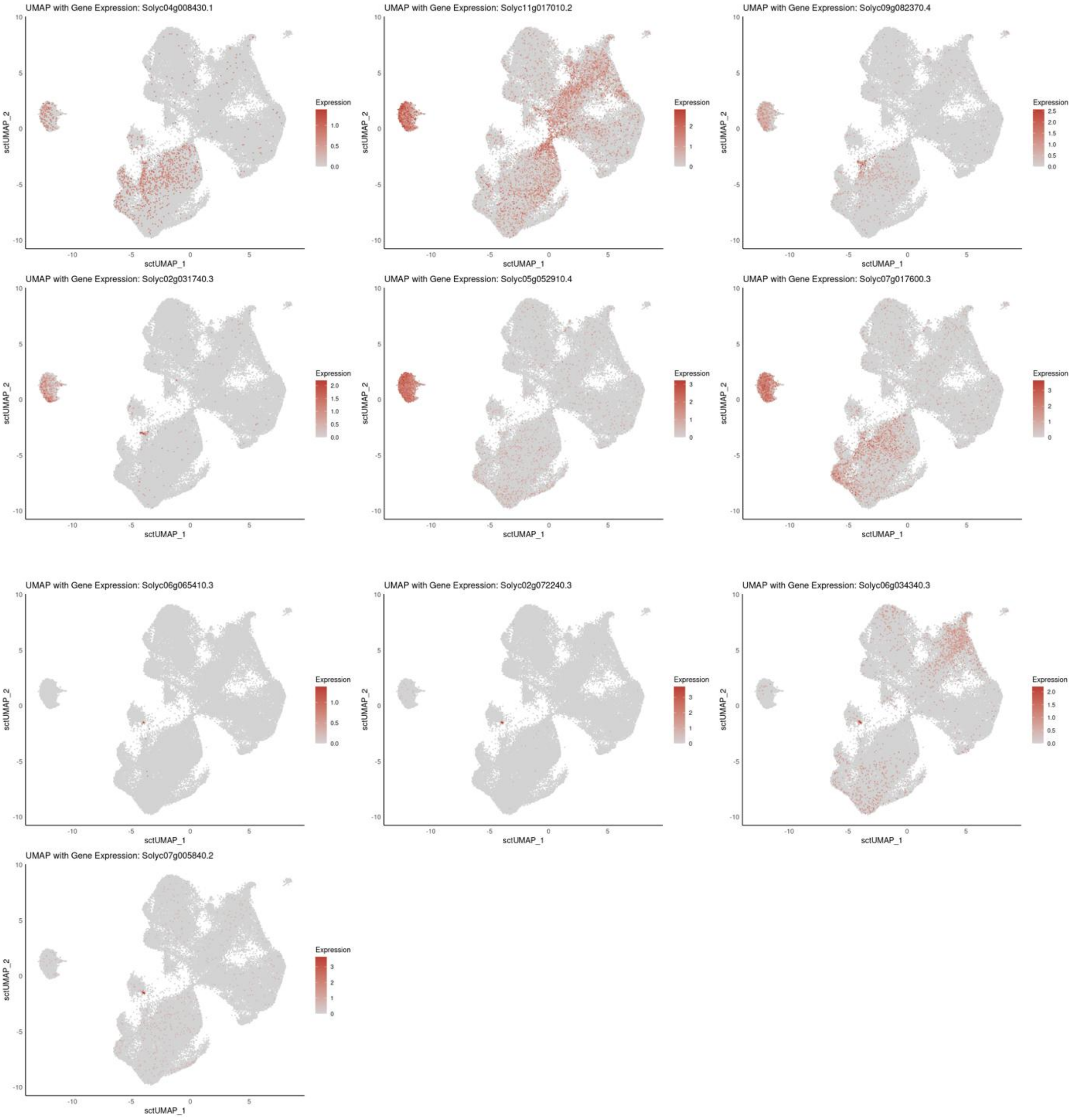


**Figure S4 (continued). The expression of individual cluster marker genes displayed on a UMAP of the snRNA-seq tomato–*R. irregularis* dataset.**

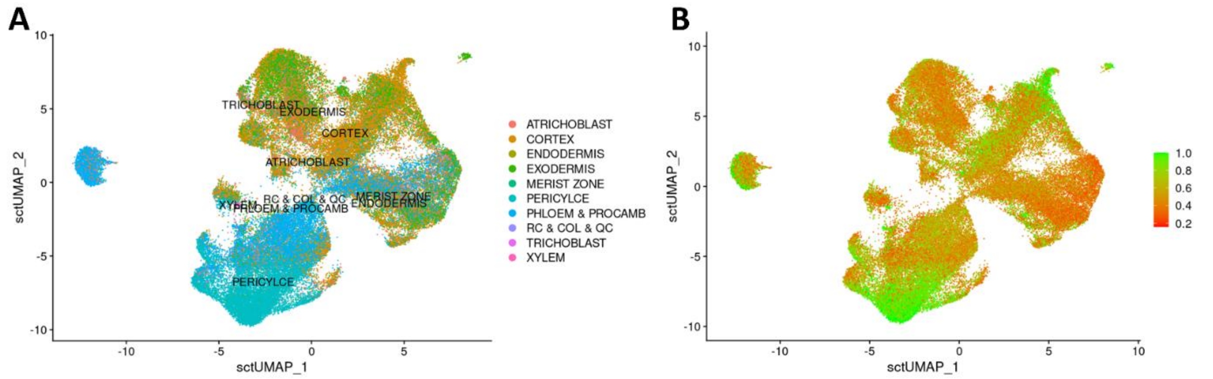


**Figure S5. Label transfer of reference tomato scRNA-seq dataset annotations onto our snRNA-seq dataset of AMF-colonized tomato roots.** A. UMAP of our tomato–AMF snRNA-seq dataset with cells colored according to the transferred cell type labels from the reference dataset of Cantó-Pastor et al. (2024). Labels were assigned using Seurat's anchor-based transfer method. B. Same UMAP colored by the prediction score for the transferred labels, reflecting the confidence of the annotation (range 0–1; higher values indicate stronger support from reference anchors).

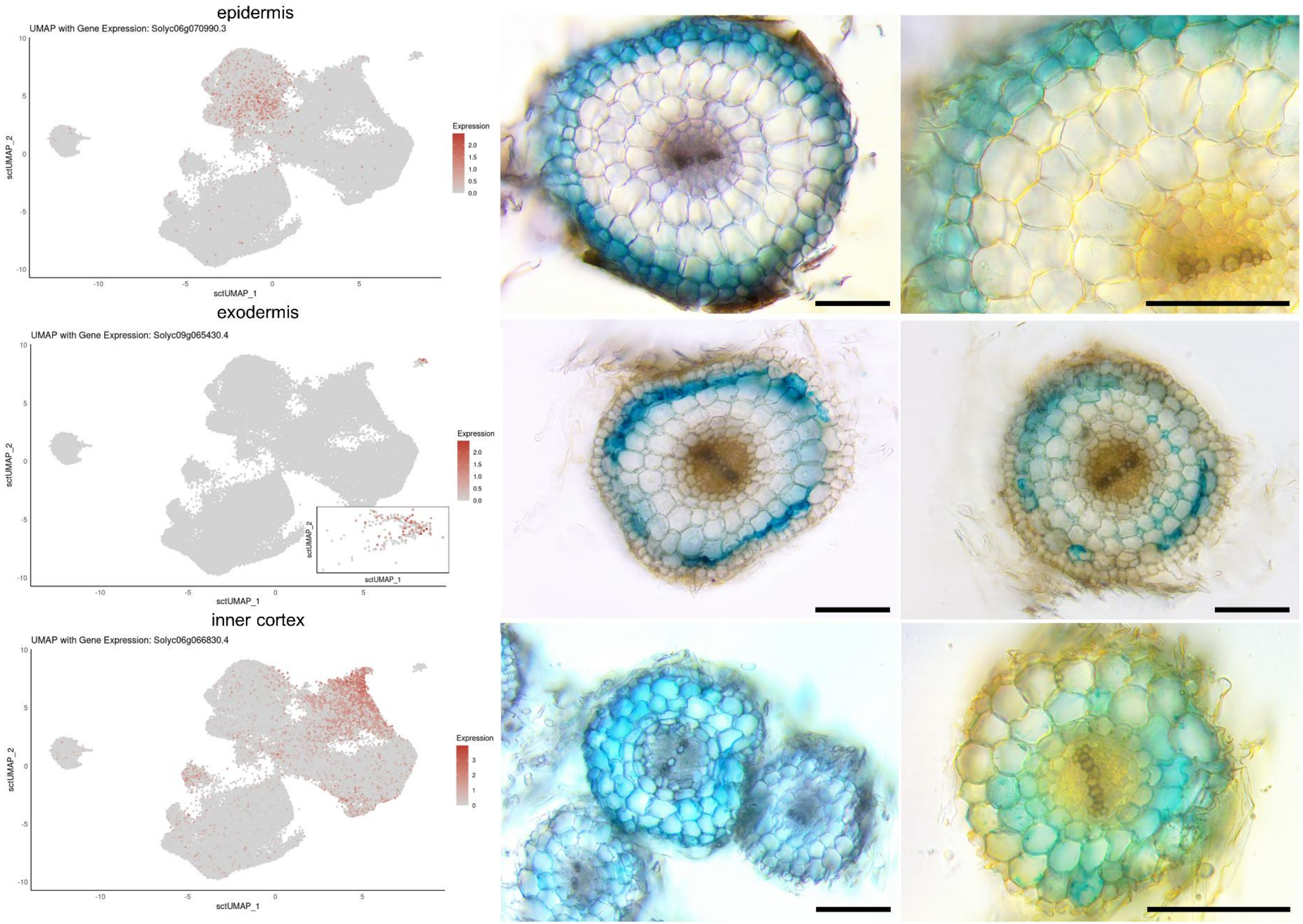


**Figure S6: Annotation of snRNA-seq clusters using promoter::GUS staining (continued on next pages).** For annotation of the snRNA-seq clusters, marker genes displaying specific expression in diverse clusters were selected, which were shown to correspond to the epidermis, exodermis, inner cortex, endodermis, phloem, and xylem. Expression profiles of marker genes are indicated on the snRNA-seq UMAP. Composite plants with roots expressing *promoter::GUS* constructs were generated using Agrobacterium-mediated hairy root transformation. GUS-stained roots were imaged using the Olympus BX51 microscope. Scale bars are 100 µm.

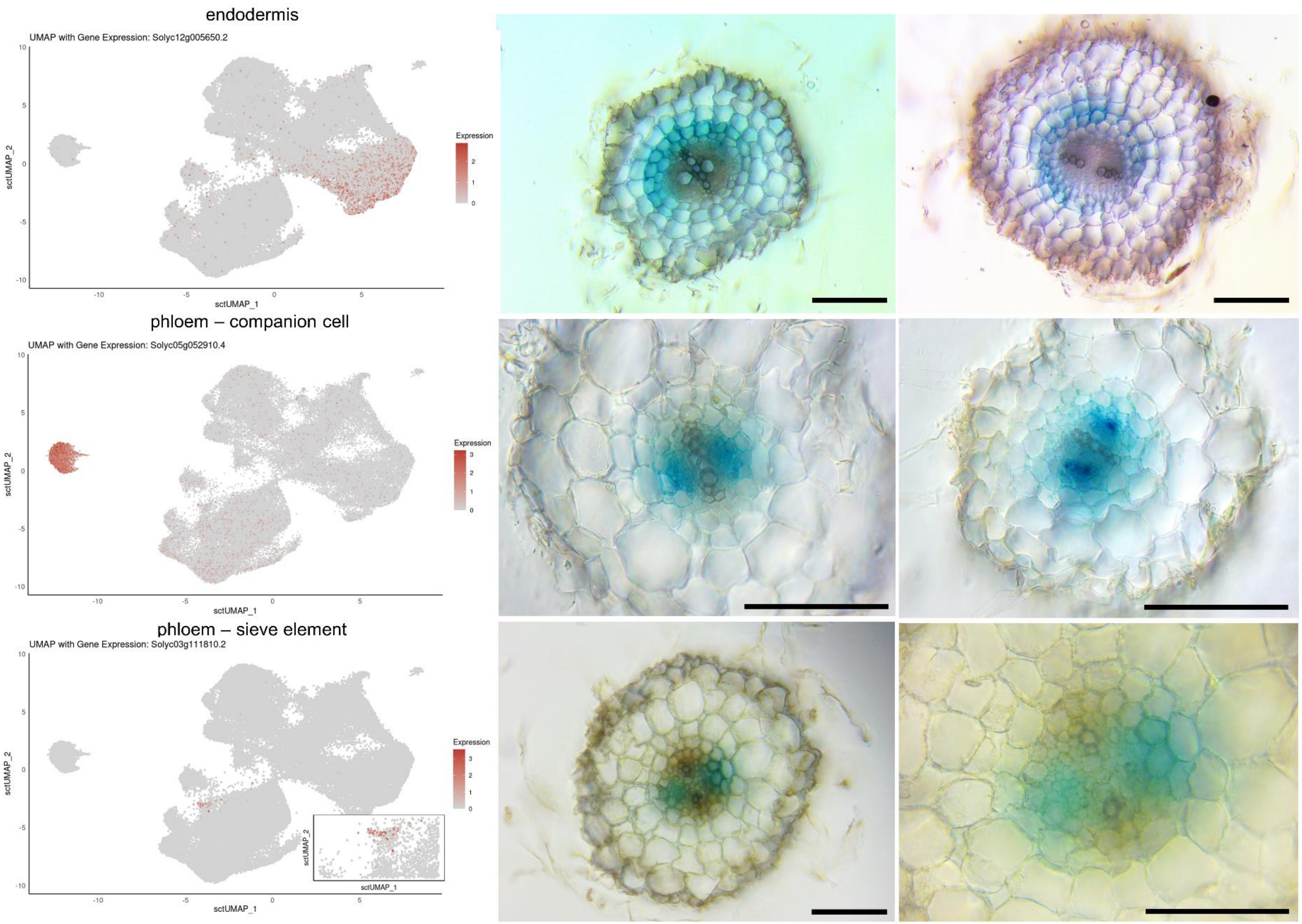
endodermis
UMAP with Gene Expression: Solyc12g005650.2
phloem – companion cell
UMAP with Gene Expression: Solyc05g052910.4
phloem – sieve element
UMAP with Gene Expression: Solyc03g111810.2
sctUMAP_1
sctUMAP_2
Expression

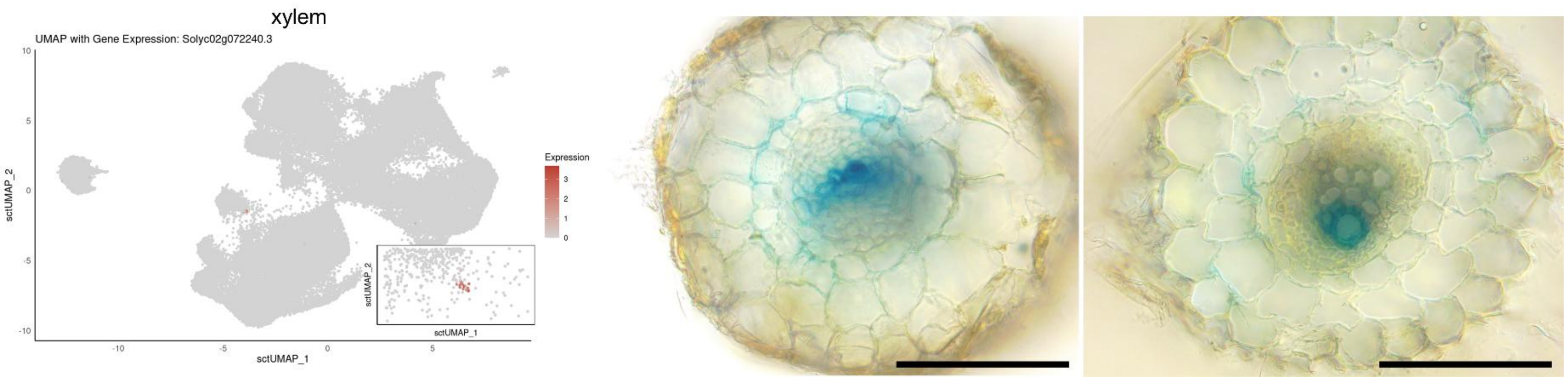
xylem
UMAP with Gene Expression: Solyc02g072240.3
sctUMAP_2
sctUMAP_1
Expression

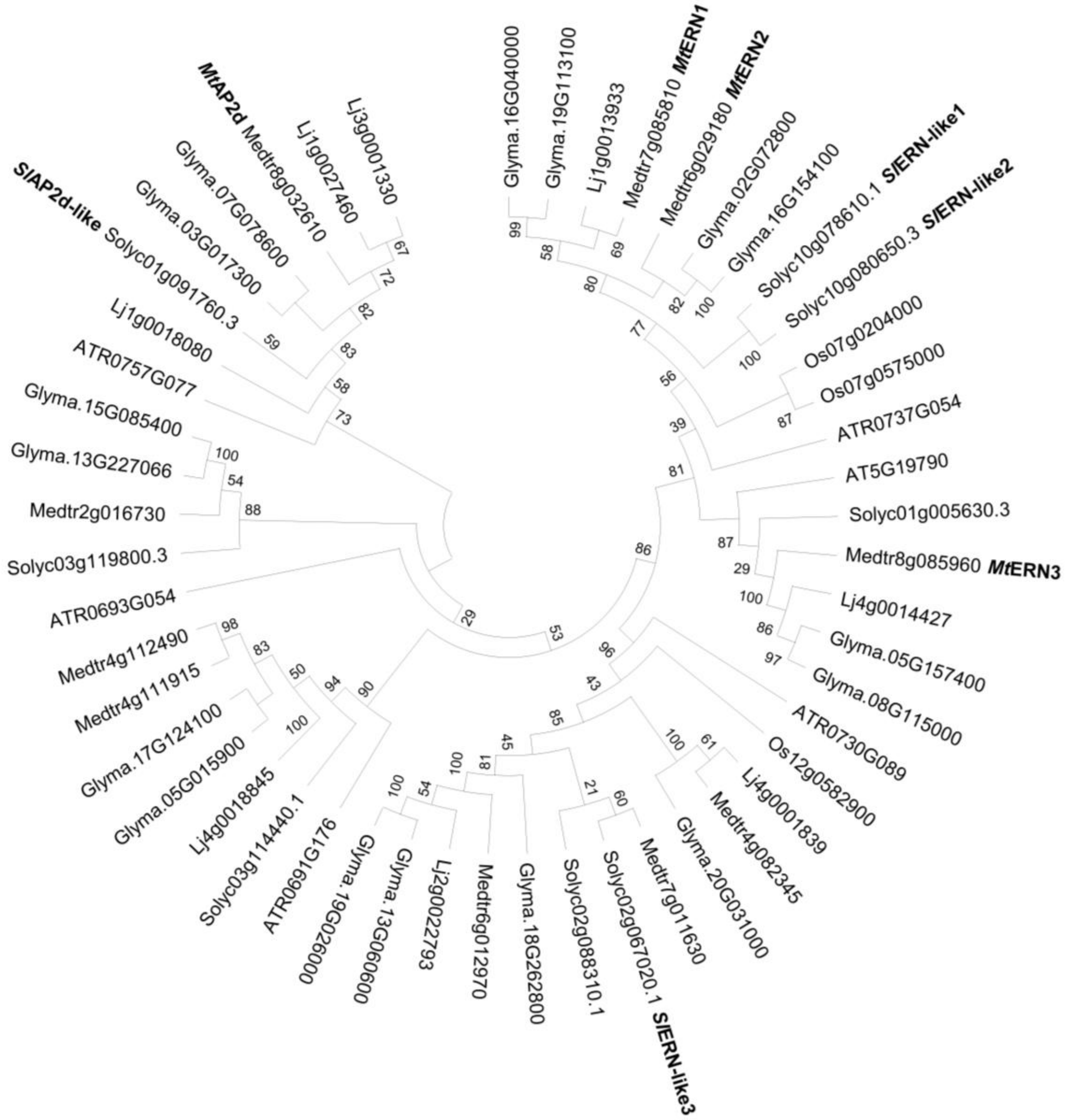


**Figure S7. Phylogenetic tree of the AP2/ERF domain homologous gene family HOM05D001032 (PLAZA5.0) encompassing ERN-like transcription factors**. The phylogenetic tree was constructed using the Maximum Likelihood method with the JTT+Freq model. The tree with the highest log-likelihood is shown. Bootstrap support values (from 117 adaptive replicates) are indicated at nodes. The initial tree for the heuristic search was selected based on log-likelihood comparison between Neighbor-Joining (NJ) and Maximum Parsimony (MP) trees. Evolutionary rate variation among sites was modeled using a discrete Gamma distribution (+G, 5 categories, shape = 0.8174) and a proportion of invariant sites (+I, 3.25%). The analysis included 51 amino acid sequences with 646 positions, and was performed in MEGA12.

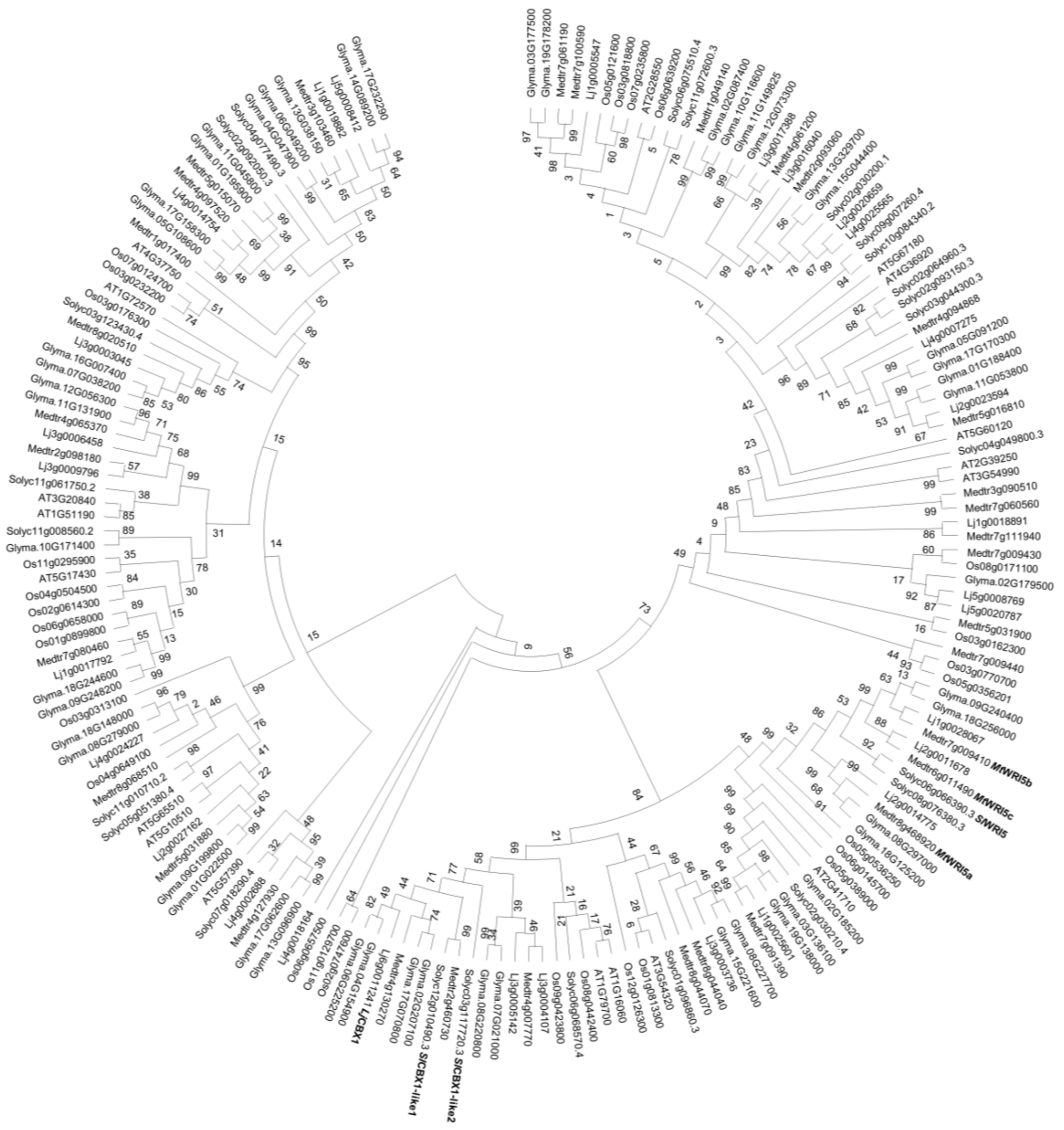

**Figure S8: Phylogenetic tree of the AP2/ERF domain homologous gene family HOM05D000138 (PLAZA5.0) encompassing *WRI5/CBX1-like* transcription factors.** The phylogenetic tree was constructed using the Maximum Likelihood method with the JTT model. The tree with the highest log-likelihood is shown. Bootstrap support values (from 128 adaptive replicates) are indicated at nodes. The initial tree for the heuristic search was selected based on log-likelihood comparison between Neighbor-Joining (NJ) and Maximum Parsimony (MP) trees. Evolutionary rate variation among sites was modeled using a discrete Gamma distribution (+G, 5 categories, shape = 0.6163) and a proportion of invariant sites (+I), which was estimated to be 0%. The analysis included 182 amino acid sequences with 1.340 positions, and was performed in MEGA12.

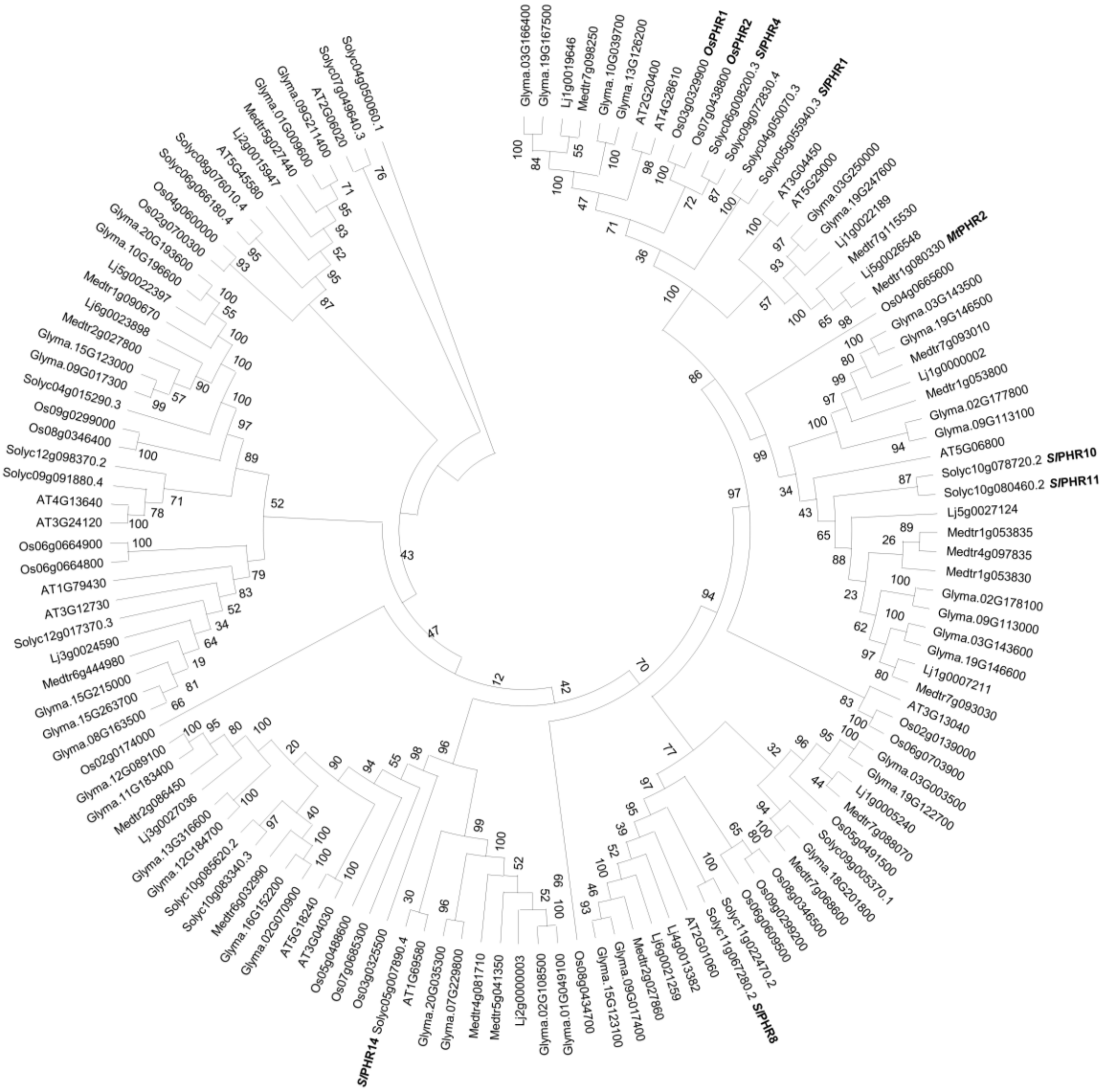


**Figure S9: Phylogenetic tree of the MYB domain PHR1-like homologous gene family HOM05D000184 (PLAZA5.0) encompassing *PHR1-like* transcription factors.** The phylogenetic tree was constructed using the Maximum Likelihood method with the JTT+Freq model. The tree with the highest log-likelihood is shown. Bootstrap support values (from 121 adaptive replicates) are indicated at nodes. The initial tree for the heuristic search was selected based on log-likelihood comparison between Neighbor-Joining (NJ) and Maximum Parsimony (MP) trees. Evolutionary rate variation among sites was modeled using a discrete Gamma distribution (+G, 5 categories, shape = 0.7901) and a proportion of invariant sites (+I, 0,12%). The analysis included 129 amino acid sequences with 860 positions, and was performed in MEGA12.

**Table S1: Overview of the cluster marker genes used for cluster validation of our snRNA-seq dataset of tomato roots**

| cell type | gene ID | gene name | Reference |
|---|---|---|---|
| Epidermis | Solyc09g090070.1 | *SlPT1* | Daram et al., (1998) |
| | Solyc09g090730.2 | *SlAMT1;1* | Filiz and Akbudak, (2020);<br>Loqué et al., (2006) |
| | Solyc12g006850.2 | *SlLKT1* | Hartje et al., (2000) |
| | Solyc10g047170.2 | *SlST1* | Howarth et al., (2003) |
| Exodermis | Solyc03g097500.3 | *SlASFT* | Cantó-Pastor et al., (2024) |
| | Solyc04g011600.4 | *SlGPAT5* | Cantó-Pastor et al., (2024) |
| | Solyc03g005320.3 | *SlKCS2b* | Cantó-Pastor et al., (2024) |
| Cortex | Solyc04g076190.1 | *SlPEP* | Kajala et al., (2021) |
| Endodermis | Solyc10g074680.2 | *SlSCR* | Kajala et al., (2021) |
| | Solyc04g077260.3 | *SlMYB36* | Li et al., (2018) |
| Pericycle | Solyc01g150101.1 | *SlPFA1* | Cantó-Pastor et al., (2024) |
| | Solyc08g008380.4 | *SlARF9B* | De Jong et al., (2015) |
| | Solyc01g068410.4 | *SlPIN5* | Mravec et al., (2009) |
| | Solyc11g011500.2 | *SlSKOR* | Cantó-Pastor et al., (2024) |
| procambium/stele | Solyc09g082830.4 | *SlAGO10* | Mirlohi et al., (2024) |
| | Solyc04g008430.1 | *SlBRL2* | Ceserani et al., (2009) |
| Phloem | Solyc11g017010.2 | *SlSUT2* | Hackel et al., (2006) |
| | Solyc09g082370.4 | *SlCVP2* | Cantó-Pastor et al., (2024) |
| | Solyc02g031740.3 | *SlPL1* | Cantó-Pastor et al., (2024) |
| | Solyc05g052910.4 | *SlLRD3* | Ingram et al., (2011) |
| | Solyc07g017600.3 | *SlPHLOEM* | Chau et al., (2023) |
| Xylem | Solyc06g065410.3 | *SlVND7* | Ramachandran et al., (2021) |
| | Solyc02g072240.3 | *SlIRX1* | Taylor et al., (2000) |
| | Solyc06g034340.3 | *SlVND6* | Kajala et al., (2021) |
| | Solyc07g005840.2 | *SlIRX3* | Taylor et al., (1999) |

**Table S2. Overview of MINI-EX–identified TFs previously linked to AM symbiosis.** Genes characterized in tomato are highlighted in pink; homologs of TFs characterized in AM symbiosis in other species, in blue; and homologs of TFs characterized in nitrogen-fixing root nodule symbiosis, in green.

<table>
<tr><th>gene ID</th><th>gene name</th><th>MINI-EX-predicted activity</th><th>described function</th><th>reference</th></tr>
<tr><td>Solyc03g123400.1</td><td>SlNSP1</td><td>Colonized epidermis</td><td>Possible ortholog of MtNSP1, which is part of the common symbiosis signaling pathway involved in early symbiotic responses</td><td>Delaux et al. (2013); Liu et al. (2011)</td></tr>
<tr><td>Solyc10g078610.1</td><td>SlERN-like1</td><td rowspan="2">Primarily in presymbiotic cortical cells and during arbuscule development</td><td rowspan="2">Close homologs of MtERN1/2/3. MtERN1/2 are involved in infection thread formation and nodule organogenesis. Upregulated during mycorrhization, but an AM symbiosis-related function is not yet described.</td><td rowspan="2">Cerri et al. (2016); Czaja et al. (2012); Kawaharada et al. (2017); Liu et al. (2023)</td></tr>
<tr><td>Solyc10g080650.3</td><td>SlERN-like2</td></tr>
<tr><td>Solyc02g067020.1</td><td>SlERN-like3</td><td>Primarily in the colonized epidermis</td><td>Close homolog of three M. truncatula AP2 TFs (Medtr6g012970, Medtr7g011630 and Medtr4g082345) AM-conserved genes, albeit of unknown function.</td><td>Bravo et al. (2016)</td></tr>
<tr><td>Solyc01g006930.3</td><td>SlNF-YA10a</td><td>Mature arbusculated cells</td><td>Close homolog of MtNF-YA1 (Medtr1g056530) and MtNF-YA2 (Medtr7g106450), involved in rhizobial infection and root nodulation.</td><td>Laloum et al. (2014)</td></tr>
<tr><td>Solyc08g007960.1</td><td>SlNF-YC3</td><td>Colonized epidermis, presymbiotic cortical cells, and cells undergoing arbuscule development</td><td>Close homolog of MtNF-YC6/MtCbf1 (Medtr2g081600) and MtNF-YC11/MtCbf2 (Medtr2g081630), AM-conserved genes with expression associated with fungal contact.</td><td>Bravo et al. (2016); Hogekamp et al. (2011)</td></tr>
<tr><td>Solyc09g066450.3</td><td>SlGRAS43</td><td>Colonized epidermis, presymbiotic cortical cells, and cells undergoing arbuscule development.</td><td>Expression of SlGRAS43 in arbusculated cells has been confirmed previously, but its RNAi-mediated knock-down did not significantly affect mycorrhizal colonization.</td><td>Ho-Plágaro et al. (2019)</td></tr>
</table>

| | | | | |
|---|---|---|---|---|
| Solyc06g009610.1 | *SlGRAS33* | Cells undergoing arbuscule development | Close homolog of *M. truncatula* Mycorrhiza Induced GRAS 1 (MtMIG1), which facilitates radial cortical cell expansion during arbuscule development. | Heck et al. (2016); Ho-Plágaro et al. (2019) |
| Solyc02g094340.1 | *SlRAM1* | Cells undergoing arbuscule development and mature arbusculated cells. | RAM1 is involved in arbuscule development and symbiotic nutrient exchange, with its role confirmed in tomato. | Ho-Plágaro et al. (2024); Luginbuehl et al. (2017); Pimprikar et al. (2016) |
| Solyc12g010490.3 | *SlCBX1-like1* | Cells undergoing arbuscule development | Close homolog of LjCBX1, which contributes to both arbuscule formation and symbiotic nutrient exchange in *L. japonicus* | Leng et al. (2023); Xue et al. (2018) |
| Solyc03g117720.3 | *SlCBX1-like2* | | | |
| Solyc06g066390.3 | *SlWRI5a* | Mature arbusculated cells | Close homolog of the *M. truncatula* WRINKLED 5a/b/c genes, which regulate nutrient exchange genes in mature arbusculated cells | Jiang et al. (2018) |
| Solyc09g005370.1 | *SlMYB1* | Mature arbusculated cells | Ortholog of MtMYB1, involved in arbuscule degradation | Floss et al. (2017) |
| Solyc10g078720.2 | *SlPHR10* | Mature arbusculated cells | Members of the PHR family with regulatory roles in AM symbiosis have been identified, including in tomato. | Das et al. (2022); Liao et al. (2022); Shi et al. (2021); Wang et al. (2024) |
| Solyc05g007890.4 | *SlPHR14* | | | |

**Table S3: Overview of the primers used in this study for domestication of marker gene promoters.**

| gene id | primer name | primer sequence | length of promoter (bp) |
|---|---|---|---|
| Solyc06g051850.2 | pSlPT4_PGGA_F | AGAAGTGAAGCTTGGTCTCAACCTTTGAAACCAGGACATATAATATTGGCAAG | 1638 |
| | pSlPT4_PGGA_R | AGGGCGAGAATTCGGTCTCATGTTCTCTAAATTGCTCAATTTACGTATATATAGAC | |
| Solyc01g096820.4 | pSlCCaMK_PGGA_F | AGAAGTGAAGCTTGGTCTCAACCTCAAATACGGCATGAGTTCAGCAC | 2950 |
| | pSlCCaMK_PGGA_R | AGGGCGAGAATTCGGTCTCATGTTTTTATTTTTTTTTGAAGAACAAAGAGAGAAAATGTTGAGTTTTGA | |
| Solyc06g070990.3 | pEPI7_PGGA_F | AGAAGTGAAGCTTGGTCTCAACCTCTCATATCGGTATAAGTGTAATTTGTGAGAC | 2947 |

| | | | |
|---|---|---|---|
| | pEPI7_PGGA_R | AGGGCGAGAATTCGGTCTCATGTTGTATAGACCTAATTATCTCCTTTTAATTTCTCTAGCTAG | |
| Solyc09g065430.4 | pEXO3_PGGA_F | AGAAGTGAAGCTTGGTCTCAACCTGTGGCTTGGAGTTAAACCTCTTTGAC | 1102 |
| | pEXO3_PGGA_R | AGGGCGAGAATTCGGTCTCATGTTTTTCTTTAAGTTTAGGGAAGTTTGAGTTTGTTGG | |
| Solyc06g066830.4 | pCOR2_PGGA_F | AGAAGTGAAGCTTGGTCTCAACCTGTTCTATATCACCATTTGCATCGTGTC | 2945 |
| | pCOR2_PGGA_R | AGGGCGAGAATTCGGTCTCATGTTATTTTCTATAAAGAGAGAACTCTCAAGTGTCTTTG | |
| Solyc12g005650.2 | pENDO2_PGGA_F | AGAAGTGAAGCTTGGTCTCAACCTGTGTGAAATATAACACCAAATAGCCTGGTG | 2931 |
| | pENDO2_PGGA_R | AGGGCGAGAATTCGGTCTCATGTTCGATCAAATGATTAATGTTCTTGATAATCCATACG | |
| Solyc05g052910.4 | pPHL3_PGGA_F | AGAAGTGAAGCTTGGTCTCAACCTTGTAAAAGAGAAATACGATAGATAACTTGAGATGGAGAG | 2668 |
| | pPHL3_PGGA_R | AGGGCGAGAATTCGGTCTCATGTTTGATATGTATAGAAACAAAAGAAATTTGTTGAGATTTTTCAAGATTTG | |
| Solyc03g111810.2 | pPHL14_PGGA_F | AGAAGTGAAGCTTGGTCTCAACCTGAGGGTCCAAAATGTTGAAATCAATCACAC | 2879 |
| | pPHL14_PGGA_R | AGGGCGAGAATTCGGTCTCATGTTTTTTTGTTGCTAATTTGGGATTTTAATTGAAGAAAAAAAAAG | |
| Solyc02g072240.3 | pXYL3_PGGA_F | AGAAGTGAAGCTTGGTCTCAACCTAACCTCTCTGAATGAAGATGTCTATTGCATG | 1211 |
| | pXYL3_PGGA_R | AGGGCGAGAATTCGGTCTCATGTTTTTTTCCACTCACAGAAACAGTGCTAGTC | |
